# Defective Viral Genomes Reveal History-dependent Self-regulation of Type I Interferon Dynamics

**DOI:** 10.64898/2026.10.30.763053

**Authors:** Yimei Li, Carla Lourens, Bjarke Frost Nielsen, Simon A. Levin, Alan S. Perelson, Bryan T. Grenfell, Aartjan J.W. te Velthuis

## Abstract

Type I interferons (IFN) help clear a viral infection, but IFN levels can then decline to reset the antiviral state and avoid innate immune dysregulation. Such dysregulation can occur following infection with highly pathogenic and pandemic RNA viruses, such as H5N1 influenza A viruses or SARS-CoV-2. In spite of the strong innate immune response that these viruses induce, pandemic viruses replicate and spread successfully, suggesting that they can circumvent the effects of IFN.

Innate immune activation is triggered when host cells detect viral RNA, especially defective viral genomes (DVGs). A strong IFN response (e.g. to DVGs or a highly immunogenic wild-type infection) must then decline through negative feedback involving the IFN receptor. How DVGs modulate this feedback remains poorly understood.

To quantify positive and negative IFN regulation, we first performed *in vitro* experiments combining different doses of viral RNA and exogenous IFN, and measured subsequent IFN expression. We next developed an *in vivo* Bayesian computational model that captures the impact of influenza virus DVGs on both positive and negative

IFN feedback. Competition between DVGs and virus is also considered. Finally, we calibrated this model on published *in vitro* and *in vivo* influenza virus infection data.

Our model suggests that the cumulative IFN exposure of a host cell, rather than the instantaneous IFN level, shapes further IFN induction; this gives DVGs a dual role: beyond triggering IFN induction, they can also lower it through negative feedback. Our model further predicts that these effects are important locally, shaping the spatial dynamics of the response, allowing suppression of innate immune responses in neighbouring, uninfected cells. Through this model, we can explain two important biological observations: i) IFN levels can decline while virus and DVG levels continue to rise in tissue culture and mouse infections; and ii) pandemic viruses that produce high levels of immunogenic RNA molecules can successfully replicate through local innate immune suppression.

Overall, we propose that heterogeneity within viral populations (including DVGs and other noncanonical viral RNAs) can modulate positive and negative feedback loops and shape infection outcomes. More broadly, our work may support studies into the therapeutic use of these viral RNA populations.

**Author summary:** Humans and animals have to fight influenza virus and other infections without inducing immunopathology. After detection of an infection, the innate immune system produces type I interferon (IFN), a signal that starts a protective response. However, high levels of IFN and prolonged IFN expression can cause damage to tissues and worsen disease. Negative feedback is one route by which IFN production can stop, but its activation during viral infection remains poorly understood.

A central question is why viruses can continue to replicate and spread despite a strong IFN response. We here investigated why IFN levels can begin to decline while virus levels continue to rise during influenza virus infection. This decline is most pronounced when the virus produces large amounts of defective viral genomes (DVGs). These defective genomes arise naturally during viral replication and are produced by many RNA viruses, including SARS-CoV-2, Ebola virus, and measles virus.

Using cell-culture experiments, computational modelling, and fitting of our model to cell-culture and mouse data, we find that defective genomes contribute to both IFN activation and its negative feedback. Both the experiments and model support a mechanism in which IFN accumulation during infection suppresses its own production, and that this suppression can start earlier under strong DVG stimulation. At the local level, this can allow viral spread to continue even while overall IFN levels remain high.

Overall, our findings reveal an additional level of interplay between defective viral genomes and the host innate immune system. They also suggest that the potential therapeutic use of defective genomes must be carefully calibrated to achieve an optimal antiviral response.

## Introduction

Influenza A viruses (IAV) cause mild to severe respiratory infections in humans and asymptomatic to deadly infections in a broad range of animal hosts. [4, 5] The clinical outcome of an IAV infection is determined in part by the virus, and by the host response to the viral infection. High inflammatory cytokine levels have been associated with immune dysregulation and high morbidity and mortality, and they occur more frequently in infections with pandemic and avian H5N1 IAV subtypes. [6–8]

Type I interferons (IFN), such as IFN-*α* and *β*, are key markers for the activation of the innate immune response during IAV infection. The production of these IFNs by infected cells helps induce an antiviral state that can reduce viral replication. [9] This process starts with the detection of viral RNA (Fig 1), including defective viral genomes, such as deletion-containing viral genomes (DelVGs), [10, 11] copy-back defective viral genomes (cbDVGs), [12] and mini viral RNAs (mvRNAs), [13, 14] by the host pathogen receptor retinoic acid-inducible gene I (RIG-I). [13, 15–17] When DVGs are packaged into particles, defective interfering particles (DIPs) are formed. [18, 19] mvRNAs, by contrast, are likely not packaged and do not spread between cells. To replicate, DVGs typically require co-infection with standard virus. [10, 18, 20, 21] In cell culture and *in vivo*, virus populations containing DVGs have been associated with stronger cytokine induction than populations that lack DVGs. [17, 22, 23] Once IFN is produced, an IFN receptor (IFNAR)-mediated positive feedback loop is maintained (Fig 2A).

**Fig 1.**
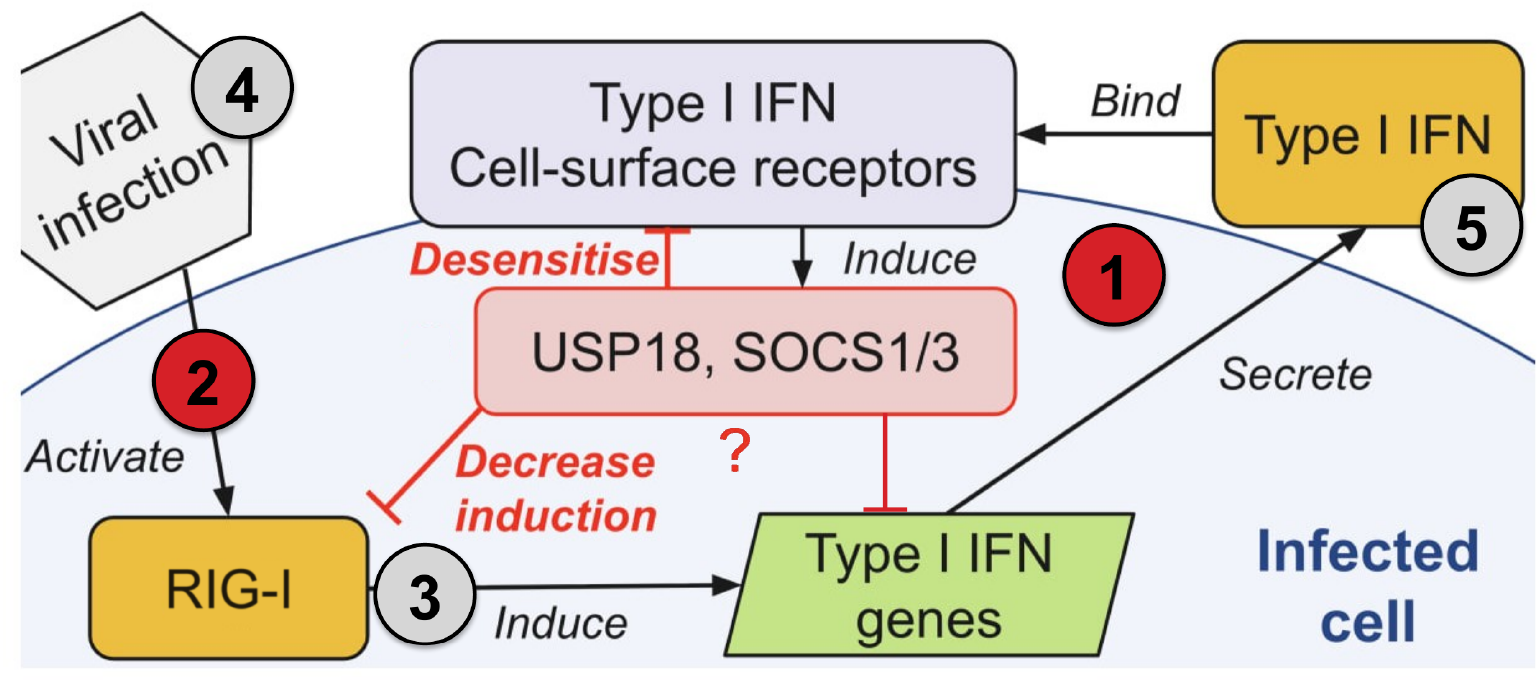
Possible sites in the type-I IFN loop where an early IFN decline can arise. Viral infection (grey hexagon) activates RIG-I (yellow rectangle, lower left), which induces type-I IFN gene expression (green parallelogram). Then, type-I IFN is secreted (yellow rectangle, upper right) and binds cell-surface type-I IFN receptors (light-purple rectangle, upper centre). Receptor signalling induces negative regulators such as USP18 and SOCS1/3 (red rectangle), which can attenuate IFN receptor signalling. Negative feedback may also act through viral RNA sensing or IFN gene expression. The red question mark indicates that the mechanism underlying the observed reduction in IFN induction remains unresolved. Numbered markers indicate five possible sites, and the Results take up each one in turn, in the order numbered here: ① IFN-driven negative feedback; ② host-mediated suppression downstream of viral RNA sensing; ③ direct antagonism of IFN induction by viral proteins; ④ loss of the inducing viral stimulus or responsive cells; and ⑤ removal or degradation of secreted IFN. Red markers indicate mechanisms retained for further consideration based on the analyses in the Results, whereas grey markers indicate mechanisms that are set aside by those analyses. Adapted from Shimoni *et al*. [1] and redrawn to include negative-feedback regulators [2, 3]. This schematic expands the feedback loop shown in Fig 2A.

**Fig 2.**
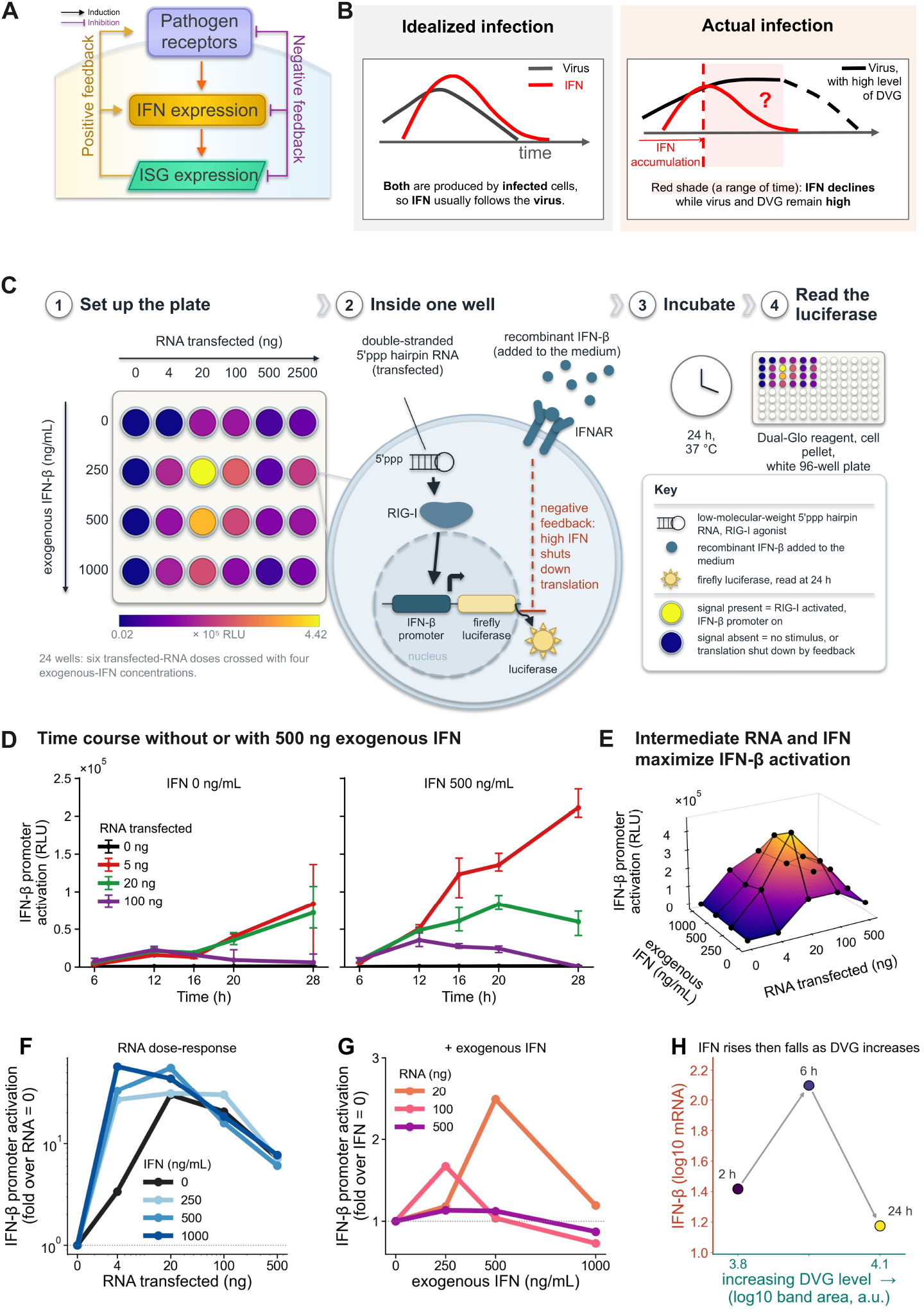
IFN dynamics and experimental measurement of IFN feedback. **(A)** Schematic of type-I IFN induction and feedback. Pathogen sensing induces IFN expression and IFN-stimulated gene (ISG) expression, with positive (yellow) and negative (purple) feedback. **(B)** Schematic of idealised virus and IFN dynamics (left) and actual IFN decline while virus with high DVG levels remains high (right). The dashed red line marks IFN accumulation. **(C)** IFN-*β* promoter–luciferase reporter assay. HEK293-Luc cells were transfected with 0–2500 ng 5^*′*^ppp hairpin RNA and exposed to 0–1000 ng/mL recombinant IFN-*β*; luciferase was measured after 24 h for the dose–response surface in (E)–(G). **(D)** IFN-*β* promoter activation at 6, 12, 16, 20, and 28 h at four RNA doses, without or with 500 ng/mL exogenous IFN. **(E)** IFN-*β* promoter activation across RNA and IFN doses at 24 h. **(F)** RNA dose–response relative to RNA = 0, at 24 h. **(G)** Exogenous IFN dose–response relative to IFN = 0, at 24 h. **(H)** IFN-*β* mRNA and DVG abundance at 2, 6, and 24 h, in the A549 infection. DVG abundance is the PB1 defective band area on the gel from Penn *et al*. Fig 3b.

However, after IFNs are produced and the viral infection is cleared, IFN levels must be reduced to limit damage. [24] Indeed, IFN exposure induces negative regulators, including ubiquitin-specific peptidase 18 (USP18), suppressor of cytokine signalling (SOCS) proteins, IFNAR regulation, and receptor internalisation (Fig 1). [2, 3, 9, 25–27] In influenza-infected respiratory epithelial cells, SOCS1 and SOCS3 induction requires a RIG-I/MAVS/IFNAR1-dependent pathway. [28] SOCS-3 can also be induced directly by viral RNA sensing through NF-*κ*B, independently of IFN and early in the infection cycle. [29] Underlining the importance of this negative feedback loop (Fig 2A), loss of type-I IFN receptor expression in mice reduces morbidity, mortality, and inflammatory cytokine levels after influenza infection. [30]

The timing of the IFN expression and its subsequent decline may affect both the viral load and the amount of tissue damage. This is also important for treatment. IFN can lose its antiviral effects and even promote inflammation once infection is established. [31] Moreover, the negative regulators weaken Janus kinase–signal transducer and activator of transcription (JAK–STAT) signalling and can make cells less responsive to repeated IFN stimulation. [3, 32] This suggests that the accumulated IFN exposure (i.e., the total IFN a cell has been exposed to during the whole infection), instead of the IFN concentration at a given time, contributes to infection outcome. Moreover, the measured systemic IFN concentration and the local, cumulative concentration that a cell has been exposed to could vary greatly, creating virus microenvironments [25, 32].

To date, within-host computational IAV models have usually included only positive IFN feedback. [33, 34] The IFN decline is then modelled by two routes: loss of extracellular IFN (written as a first-order clearance term, together with receptor binding) and a fall in IFN induction (less IFN secreted by infected cells). [33–35] The latter factor has been coupled to infected-cell abundance, which decreases over time through several mechanisms: target-cell depletion, infection, and IFN-induced refractoriness. [2, 34, 36] These assumptions tie IFN induction to infected-cell abundance, so IFN induction only starts to decline once infected-cell abundance falls. As a result, the decline in IFN levels can lag behind the decline in viral levels (Fig 2B left). However, several *in vitro* and *in vivo* studies show the opposite behaviour: IFN declines while infectious virus or viral RNA levels stay high (Fig 2B, right; S1 Fig). [20, 37, 38] A live attenuated influenza vaccine containing DIPs induced less type-I IFN and showed delayed but improved replication in mice, so more virus did not mean more IFN. [39] Moreover, pandemic and highly pathogenic avian IAV strains induce high IFN and other cytokine levels, while successfully growing to high titres and spreading. [6–8] Finally, while target cell levels can be depleted in tissue culture, their depletion is less likely to occur at the scale of a whole organ or organism. [33, 34, 40–42]

We here asked whether explicit negative feedback from past IFN, a term whole-host models currently do not consider, [33, 34, 40] could better describe the interactions between the viral infection and the host immune system. Moreover, we hypothesise that the viral RNA molecules that activate IFN expression should also be considered as factors that can, through the type-I IFN receptor (IFNAR), reduce IFN expression. Experimentally, we confirm that titration of a synthetic viral-RNA mimic (RIG-I ligand) in the context of exogenous type-I IFN jointly and nonlinearly shapes further IFN induction. To model this, we introduce negative feedback on IFN induction as a regulatory force. Using *in vivo* and A549 cell data, our model can reproduce the early IFN decline in high-DVG conditions. For broader impact, at a local level, our spatial simulation predicts that negative feedback enables virus spread, even when IFN levels are high overall, allowing continued viral spread. At the intervention level, we show that the outcomes of DVG- and IFN-based treatments depend on host state and treatment timing. Overall, our findings are important for understanding influenza dynamics, host-pathogen immune interactions, and DVG- and IFN-based interventions in general, such as therapeutic interfering particles (TIPs). [43, 44] The paper follows this order: the reporter experiment first, then the whole-host model fitted to the mouse time series, and then two directions that follow from it — the spatial consequences of local feedback, building on our earlier *in vitro* spatial model [45], and the implications for DVG- and IFN-based treatment.

## Results

### Result 1: IFN and immunogenic RNA jointly regulate IFN induction

An early decline in measured IFN during infection can be explained by many different mechanisms, and whole-host time courses of virus, DVG, and IFN do not separate them. (Possible explanations act at different places in the IFN loop, Fig 1: ① IFN-driven negative feedback, ② host-mediated suppression downstream of viral RNA sensing, ③ direct antagonism of IFN induction by viral proteins, ④ loss of the inducing viral stimulus or responsive cells, and ⑤ removal or degradation of secreted IFN.) One recurring difficulty in within-host viral dynamics modelling is distinguishing immune regulation from changes in target-cell availability, because both can produce similar viral trajectories [34, 46]. For example, low IFN levels can permit continued viral replication, whereas cells in antiviral states induced by high IFN levels may reduce infection but preserve susceptible cells for later rounds of spread. [46] To systematically test how viral RNA stimulation and exogenous IFN affect the decline in IFN-*β* promoter activity, we transfected a titration series of a synthetic RIG-I ligand in the presence or absence of recombinant IFN-*β* into HEK293-Luc cells (Fig 2C) [47, 48]. Because this assay uses noninfectious 5^*′*^ppp hairpin RNA rather than infectious virus, it excludes viral replication, viral protein expression, and cell-to-cell spread, so changes in target-cell availability and viral antagonism cannot explain the IFN reporter dynamics. In addition, variation of the exogenous IFN level allowed us to ask whether the history of IFN exposure affects when further IFN induction begins to decline. Finally, since our readout is IFN-*β* promoter activation rather than IFN protein level, the removal or degradation of secreted IFN is not a confounding factor in asking whether the level and timing of IFN-*β* induction itself can change under different combinations of RNA stimulation and extracellular IFN. An assay without infectious virus tests this regulation directly (Box 1).

In the time-course experiment, we observed a decline in IFN promoter activity at 100 ng RNA transfected in the absence of exogenous IFN, but not at lower RNA levels (Fig 2D). This behaviour is in agreement with previous reporter experiments showing a strong increase in IFN-*β* promoter activity with increasing amounts of transfected immunostimulatory RNA [45, 47]. In contrast, when exogenous IFN was added, the maximum IFN promoter activity increased, but the decline in IFN promoter activity also occurred at lower RNA concentrations, for example at 20 ng of RNA with 500 ng/mL exogenous IFN, compared with 100 ng of RNA without exogenous IFN (Fig 2D). Thus, increasing the RNA stimulus affected not only the magnitude of IFN-*β* induction but also the timing of its peak and subsequent decline. To better reveal this interplay, we performed RNA and IFN titrations and identified a peak of IFN-*β* induction at 20 ng of RNA and 250 ng/mL exogenous IFN (Fig 2C and E).

Moreover, we observed that high exogenous IFN levels can trigger suppression of IFN promoter activity at lower RNA levels (Fig 2F), whereas high RNA levels can suppress IFN promoter activity (Fig 2G). This pattern is consistent with cumulative-IFN feedback, in which the history of IFN exposure progressively reduces the capacity for further IFN induction, rather than IFN acting instantaneously.

Because the assay excludes viral replication, viral proteins, and secreted IFN, three candidate mechanisms are set aside (Fig 1), leaving two: ① IFN-driven negative feedback and ② host-mediated suppression downstream of viral RNA sensing. Our results support these two acting together to shape subsequent IFN-*β* induction.

The observed RNA and IFN dose dependence suggests that DVG production could change not only the magnitude of IFN promoter induction, but also when that induction begins to decline. In line with the transfection results, a similar decline in IFN-*β* expression was observed during IAV infection *in vitro* using a virus population that contained DVGs (Fig 2H and S2 Fig). Together, these experiments support a role for IFN exposure in the decline of subsequent IFN induction: when exogenous IFN was added, the decline in IFN-*β* promoter activity occurred at lower RNA doses, consistent with a contribution from cumulative IFN exposure.

#### Box 1. Reporter experiment design: isolating RNA × IFN effects on subsequent IFN induction (Fig 2C)

**Time-series experiment**. HEK293-Luc cells carry firefly luciferase downstream of the IFN-*β* promoter. Cells received a 5^*′*^-triphosphate (5^*′*^ppp) double-stranded hairpin RNA acting as a RIG-I ligand, with or without 500 ng/mL recombinant IFN-*β*. Promoter activity was followed from 6 to 28 h across four RNA doses (Fig 2D).

**Design: RNA***×* **IFN dose–response surface**. At 24 h, RNA doses from 0–2500 ng were crossed with exogenous IFN from 0–1000 ng/mL, allowing the joint effect of RNA and IFN on subsequent IFN induction to be mapped (Fig 2F, G).

**Mechanistic specificity**. This 5^*′*^ppp hairpin RNA is a synthetic RIG-I agonist (not a full-length viral genome): it is noninfectious and does not spread between cells. This removes viral replication, viral proteins, and infection-driven target-cell depletion. The readout is IFN-*β* promoter activation rather than secreted IFN protein, separating induction from IFN clearance and degradation.

**Main result**. RNA and IFN jointly regulate subsequent IFN induction nonlinearly. In the time series, exogenous IFN shifted the decline in promoter activity to lower RNA doses (Fig 2D). Across the 24 h response surface, increasing IFN shifted suppression toward lower RNA stimulation, while high RNA also reduced induction across IFN conditions (Fig 2F, G).

**Conclusion**. By removing viral replication, viral proteins, target-cell depletion, and IFN clearance, this experiment narrows the possible causes of reduced IFN induction to two host-mediated mechanisms: IFN negative feedback and RNA-triggered suppression (Fig 1).

**Limitations**. The absence of virus is both the strength and the limit of this design. Without replication, viral antagonists, or cell-to-cell spread, the assay cannot report how those processes feed back on IFN induction during an infection. The single step dose of exogenous IFN is also not the gradual accumulation a cell experiences *in vivo*.

### Result 2: A model with cumulative-IFN driven negative feedback

The reporter experiment revealed an unexpected pattern: IFN induction declined even while the RNA stimulus remained high or continued to rise (Fig 2F and G). The same pattern appears in published infectious-virus datasets with high levels of DVGs or other non-canonical viral genomes (Fig 2H), including IAV, [20, 37, 39] respiratory syncytial virus, [49, 50] and Sendai virus, [51] across both *in vitro* and *in vivo* systems (project repository, GitHub). Among these datasets, Penn *et al*. [20] provide a particularly informative *in vivo* IAV time series, with longitudinal measurements of IFN, virus, and DVG abundance (Fig 3A). We used this dataset to test whether a model incorporating both positive and negative IFN regulation could capture the observed IFN dynamics.

**Fig 3.**
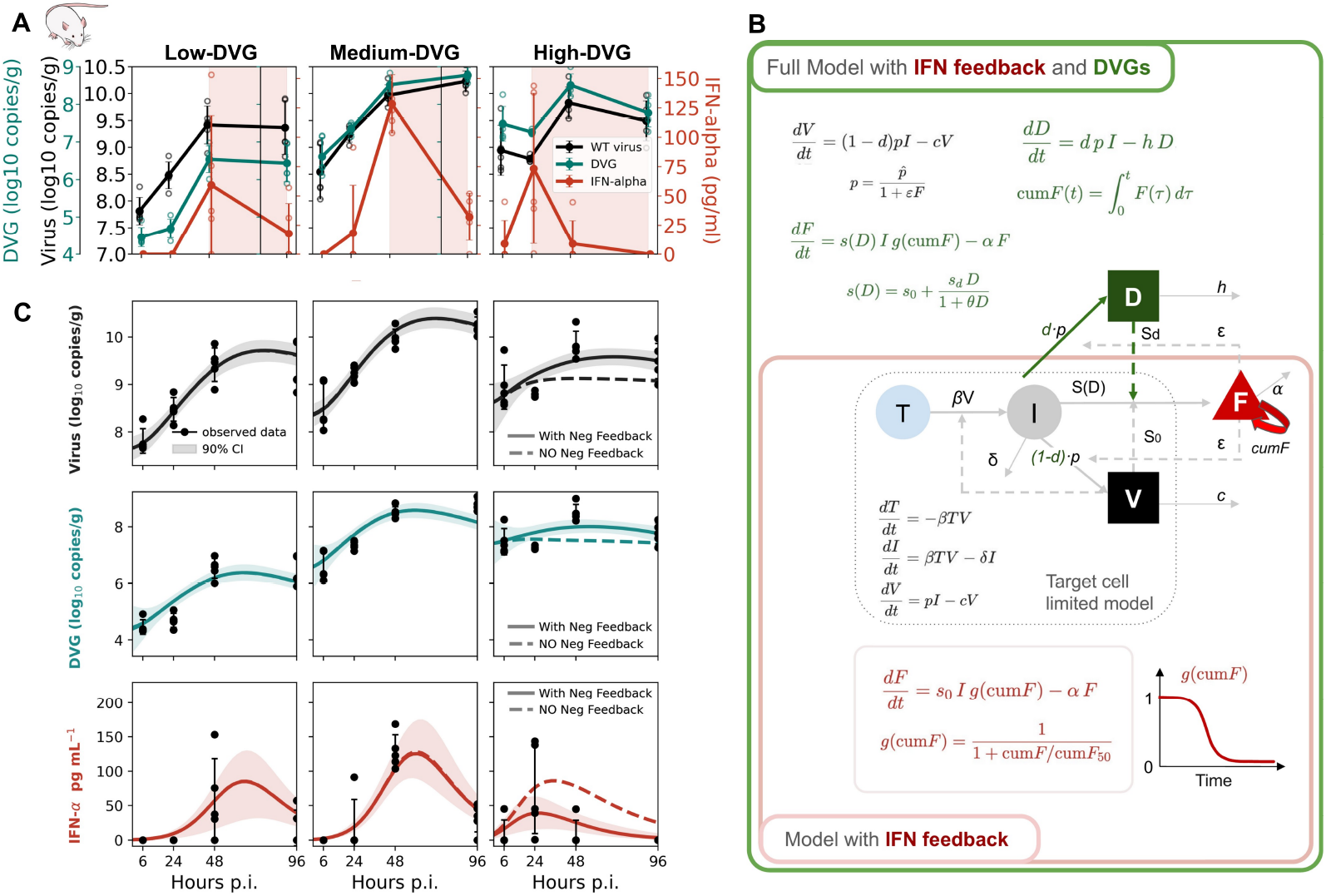
In vivo influenza dynamics, within-host model, and model fits across DVG conditions. **(A)** *In vivo* virus, DVG, and IFN-*α* measurements in low-, medium-, and high-DVG inoculum groups [20]. Virus and DVG are read from their Fig 5f and IFN-*α* from their Fig 6b, with the DVG fractions taken from their Table S1. These groups are the 6:2 Tky/05, 7:1 Tky/05 LOW, and 7:1 Tky/05 HIGH stocks of Penn *et al*., respectively, all given at 10^5^ PFU. Open circles show individual mice, filled circles show group means, and bars show standard deviations. Shaded regions indicate the period of IFN decline. **(B)** Within-host model incorporating target cells, infected cells, virus, DVGs, and cumulative IFN negative feedback. The target-cell–virus–IFN backbone (innermost boxes) follows Baccam *et al*. [40]. Model equations and parameters are given in Eqs. 1–3 and Table 1; model variants are described in Methods. **(C)** Model fits to virus, DVG, and IFN-*α* across the three DVG conditions. Solid curves show the model with cumulative IFN feedback; dashed curves show the corresponding trajectories with negative feedback removed. Shading shows the 90% posterior interval and points show the data. (Mouse drawing: NIAID NIH BioArt Source, BIOART-000283, public domain.)

Previous observations (Fig 3A) and our reporter measurements (Fig 2C–E) indicate three features of IFN negative feedback: dependence on IFN exposure, history, and dependence on immune-activating RNA. Based on these data, we developed a simple whole-host ordinary differential equation (ODE) model with both positive and negative IFN feedback (Fig 3B), and we extended the system with DVG levels to fit the DVG-resolved data.

**Table 1.** Quick reference: symbols in the DVG-extended ODEs (Fig 3B).

| Symbol | Meaning |
|---|---|
| *T*, *I*, *V*, *D*, *F*, cum*F* | Target cells; infected cells; full-length viral RNA; DVGs; IFN; cumulative IFN exposure. |
| *β*<sup>\*</sup> | Infection rate (pairs with *T* and *V*). |
| *δ*<sup>\*</sup> | Infected-cell loss rate. |
| *p̂*<sup>\*</sup> | Baseline viral output per infected cell per day. |
| *ε*<sup>\*</sup> | How strongly IFN lowers viral output, via *p*(*F*) = *p̂*/(1 + *εF*). |
| *c*<sup>\*</sup>, *h*<sup>\*</sup> | Clearance of *V* and *D*. |
| *d*<sup>\*</sup> | Share of output that is DVGs, 0 < *d* < 1; infectious share is 1 − *d*. |
| *s*<sub>0</sub><sup>\*</sup>, *s*<sub>*d*</sub><sup>\*</sup>, *θ*<sup>\*</sup> | Baseline IFN induction; DVG boost; saturation scale in *s*. |
| *α*<sup>\*</sup> | IFN removal rate. |
| cum*F*<sub>50</sub><sup>\*</sup> | Cumulative IFN exposure at which the IFN source is reduced by half. |
| *T*<sub>0</sub>, *V*<sub>0</sub><sup>\*</sup>, *D*<sub>0</sub><sup>\*</sup>, *F*<sub>0</sub> | Initial target cells; initial virus; initial DVGs for with-DVG models; initial IFN, fixed at 10<sup>−10</sup> as numerical zero. |

<sup>\*</sup> Fitted parameters; the three groups are fitted jointly, some parameters taking one value across groups and others free or partially pooled (S2 Table). Fitting results can be found in S1 Table.

#### A whole-host model with DVGs and both positive and negative IFN feedback

Our model builds on the target-cell–virus–IFN framework introduced for IAV by Baccam *et al*. [40] and used in [33, 34, 41]. The model simply tracks target cells *T*, infected cells *I*, virus *V*, DVGs *D*, type-I IFN *F*, and the cumulative IFN exposure cum*F* (Fig 3B; symbols are summarised in Table 1). In this paper *V* represents full-length viral RNA rather than infectious virus. It is fitted to the M-gene signal with the defective share removed (Methods); the plaque-titre fits are on GitHub. Infected cells *I* produce progeny at rate *p*, and a fraction *d* ∈ [0, 1] of this progeny consists of DVGs (*D*), while the remaining (1 − *d*) is virus (*V* ):

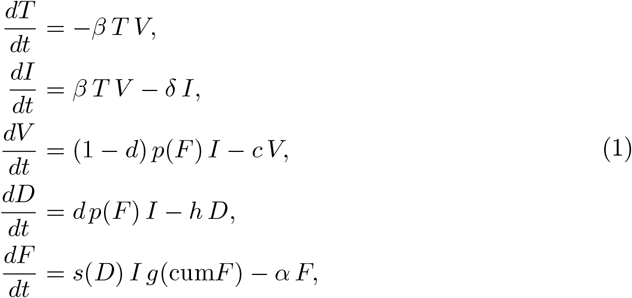

with initial conditions *T* (0) = *T*_0_, *I*(0) = 0, *V* (0) = *V*_0_, *D*(0) = *D*_0_, *F* (0) = 0, cum*F* (0) = 0. DVGs are cleared at rate *h* and modulate IFN induction through the saturation function

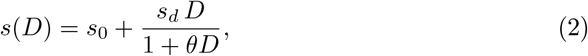

representing DVG sensing, which approaches the limit *s*_0_ + *s*_*d*_*/θ* as *D* grows; *s*_0_ is the baseline IFN induction rate without DVGs. All other rates are inherited from the core model, including the IFN-suppressed production rate 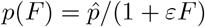 and the cumulative feedback *g*(cum*F* ) (Eq. 3). Thus, setting *d* = 0 with *D*(0) = 0 recovers the core model (Eq. 4, in Methods). The DVG-extended within-host structures follow [52–55], whereas the DVG–IFN sensing link follows [56].

The cumulative-exposure factor and the cumulative IFN exposure are

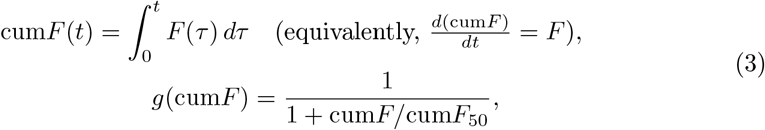

where cum*F* (*t*) is the total IFN the host system has been exposed to up to time *t* and cum*F*_50_ is the value at which the induction source is reduced by half (Fig 3B). The function *g*(cum*F* ) is a decreasing multiplier on IFN induction; it modulates the source term rather than adding a separate *F* -dependent loss term to *dF/dt*. The biological motivation for cumulative-IFN feedback follows the USP18-associated desensitisation mechanism described by [3, 57]. Because this feedback responds to the amount of IFN accumulated rather than to the specific stimulus, it is a general property of the IFN system; we hope this generality makes the framework useful in other IFN-inducing settings (see Discussion).

In order to test this model we fit it to data derived from an experiment performed by Penn *et al*. [20] in which influenza viral stocks with differing levels of DVGs were generated, here called low, medium and high. These stocks were used to intranasally infect 6–8 week old female BALB/c mice with 10^5^ PFU of virus. The model fits were done using Stan as detailed in Methods.

#### Current vs cumulative IFN feedback

We first asked whether negative feedback is needed at all by setting *g*(cum*F* ) ≡1, so that there is no effect of cumulative IFN exposure on IFN induction. Without this negative feedback, the model cannot reproduce the pronounced IFN decline in the high-DVG group and shifts the IFN peak substantially later than observed (Fig 3C, dashed). Profile likelihood further shows that feedback is supported primarily by the high-DVG data, whereas the low- and medium-DVG groups do not require it (Methods). This is also supported by our experiment (Fig 2).

We next asked what signal drives this feedback: current IFN levels or accumulated IFN exposure. The two mechanisms make different predictions: feedback driven by current IFN levels weakens as IFN levels fall, whereas cumulative IFN exposure continues to increase and can therefore sustain suppression of further IFN induction. We tested this distinction by comparing two feedback forms that differ only in the signal they respond to: *g*(cum*F* )=1/(1+cum*F* /cum*F*_50_) and *g*(*F* ) = 1*/*(1 + *F/F*_50_). The two forms are difficult to distinguish at low and medium DVG, whereas the cumulative form is better supported at high DVG, where IFN declines sharply while virus remains high (Supporting Information). Consistent with this result, the cumulative IFN-based feedback threshold (cum*F*_50_) is constrained by the high-DVG data, but not by the low- and medium-DVG data. Similar results are obtained when the DVG compartment is removed or under alternative model formulations (Materials and Methods; Supporting Information). At a molecular level, cumulative IFN-based feedback could be established in part by post-translational modifications and gene inactivation, in line with published findings. [2, 3, 25]

An alternative explanation for the IFN level decline in vivo is faster IFN removal rather than reduced production. However, faster IFN clearance cannot explain the early timing of the IFN decline, particularly its magnitude in the high-DVG group. Increasing the IFN clearance rate (*α*) within the range supported by published within-host estimates [33, 34] does not rescue the model fit, regardless of whether *α* is shared across DVG groups or fitted separately (Supporting Information, Methods, Discussion). A mathematical analysis, consistent with our reporter experiment, further shows that positive IFN regulation alone cannot reproduce the decline while the viral stimulus continues to increase (Supporting Information of Analytic argument). Thus, neither faster IFN removal nor positive regulation alone accounts for the observed IFN decline.

Taken together, our fits and analytic results support a framework combining positive IFN induction with negative feedback.

#### DVGs alter IFN expression through direct sensing and changes in the infected-cell population

We next asked why increasing DVG production does not necessarily increase IFN. Because DVGs are strong stimulators of IFN, a natural expectation is that more DVG production should produce more IFN. However, DVGs affect IFN through two routes: direct DVG sensing increases IFN induction per infected cell, whereas competition with virus reduces infectious progeny and can therefore reduce the infected-cell population. We varied the DVG production fraction (*d*) within each fitted DVG condition while keeping the initial DVG abundance (*D*_0_) fixed. These parameters describe different stages of infection: *D*_0_ represents the incoming DVG inoculum, whereas *d* determines how newly produced progeny are divided between virus and DVGs. We then followed cumulative IFN exposure, negative feedback, effective IFN induction, and the infected-cell population (Fig 4, Rows 1–4).

**Fig 4.**
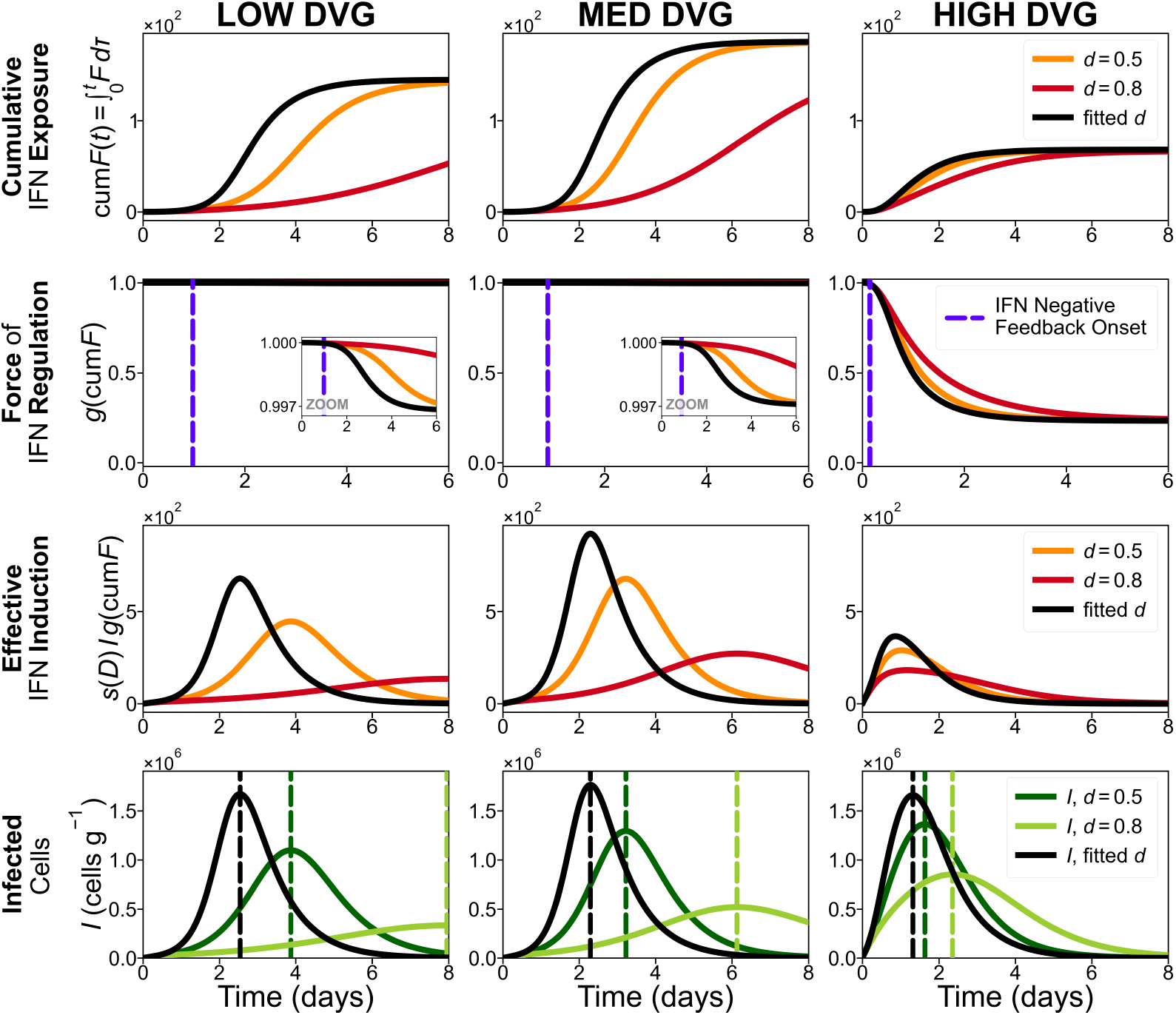
IFN regulation across DVG production fractions. Columns show the low-, medium-, and high-DVG conditions. Coloured curves show DVG production fractions (*d* = 0.5 and 0.8), and black curves show the fitted condition, whose posterior median *d* is 0.0034, 0.105 and 0.182 in the low-, medium- and high-DVG groups (credible intervals in S1 Table). Row 1 shows cumulative IFN exposure (cum*F* ). Row 2 shows the IFN-regulation term (*g*(cum*F* )); a dashed vertical marks feedback onset for each curve, taken as the time its own decline has run one per cent of its way, and insets magnify the low- and medium-DVG conditions. Row 3 shows effective IFN induction (*s*(*D*)*Ig*(cum*F* )). Row 4 shows the infected-cell population (*I*, cells g^*−*1^), with a dashed vertical at each curve’s own peak.

Unexpectedly, increasing the DVG production fraction *d* reduced cumulative IFN exposure (Row 1). Across these trajectories, DVG-dependent induction was already near saturation. Increasing *d* therefore added little through direct DVG sensing, but shifted progeny away from virus and reduced the infected-cell population (Row 4). This reduction in infected cells lowered IFN production and was most apparent in the low- and medium-DVG conditions. In these groups, *g*(cum*F* ) changed little, so negative feedback contributed little to the differences in IFN (Row 2).

The high-DVG condition revealed an additional effect of negative feedback. Here, *g*(cum*F* ) declined substantially, indicating strong suppression of further IFN induction (Row 2). Increasing *d* still reduced the infected-cell population and cumulative IFN exposure, but the lower cumulative exposure also weakened negative feedback, leaving *g*(cum*F* ) higher. Thus, higher DVG production had two opposing effects: fewer infected cells reduced IFN production, while weaker negative feedback partially restored the capacity for IFN induction.

Effective IFN induction, *s*(*D*)*Ig*(cum*F* ), shows which effect dominated (Row 3). In the high-DVG condition, induction rose early with the rapid expansion of infected cells and then declined as cumulative IFN increased and negative feedback strengthened.

This decline began before the infected-cell population itself peaked. Across all three DVG conditions, increasing *d* reduced and delayed effective IFN induction. Ultimately, for this mouse data, the reduction in the infected-cell population outweighed both additional DVG sensing and the accompanying weakening of negative feedback.

### Result 3: Local negative IFN feedback supports viral spread despite strong host-level responses

Our whole-host results revealed strong negative feedback in IFN production through systemic IFN levels and modulation by DVGs. However, upon infection, it takes time for IFN levels to rise and trigger negative feedback. In the intervening time, the antiviral response could limit or prevent viral growth. Since seasonal and pandemic IAV strains can grow to high titres in spite of producing aberrant RNA molecules and activating the immune response early in infection, how do these viruses replicate and spread despite inducing a strong antiviral response? [13, 20]

IFN exposure is expected to be highest near an infected focus and decrease with distance (Fig 5, top left). [45, 58–60] However, recent work has revealed that the antiviral response is not a simple gradient and that cells directly adjacent to the infected focus (i.e., neighbouring cells) are more susceptible to infection than cells located more distally from the focus. [61] Since the mechanism underlying this ordered structure is not fully understood, we wondered if viruses could hijack the negative feedback system on a local scale and establish a microenvironment that would enable them to replicate and spread. Our earlier *in vitro* spatial model of IFN spread is the starting point here [45]; there the local IFN response was monotonic in distance, and we now ask what changes when the reporter’s non-monotonic response is used instead.

**Fig 5.**
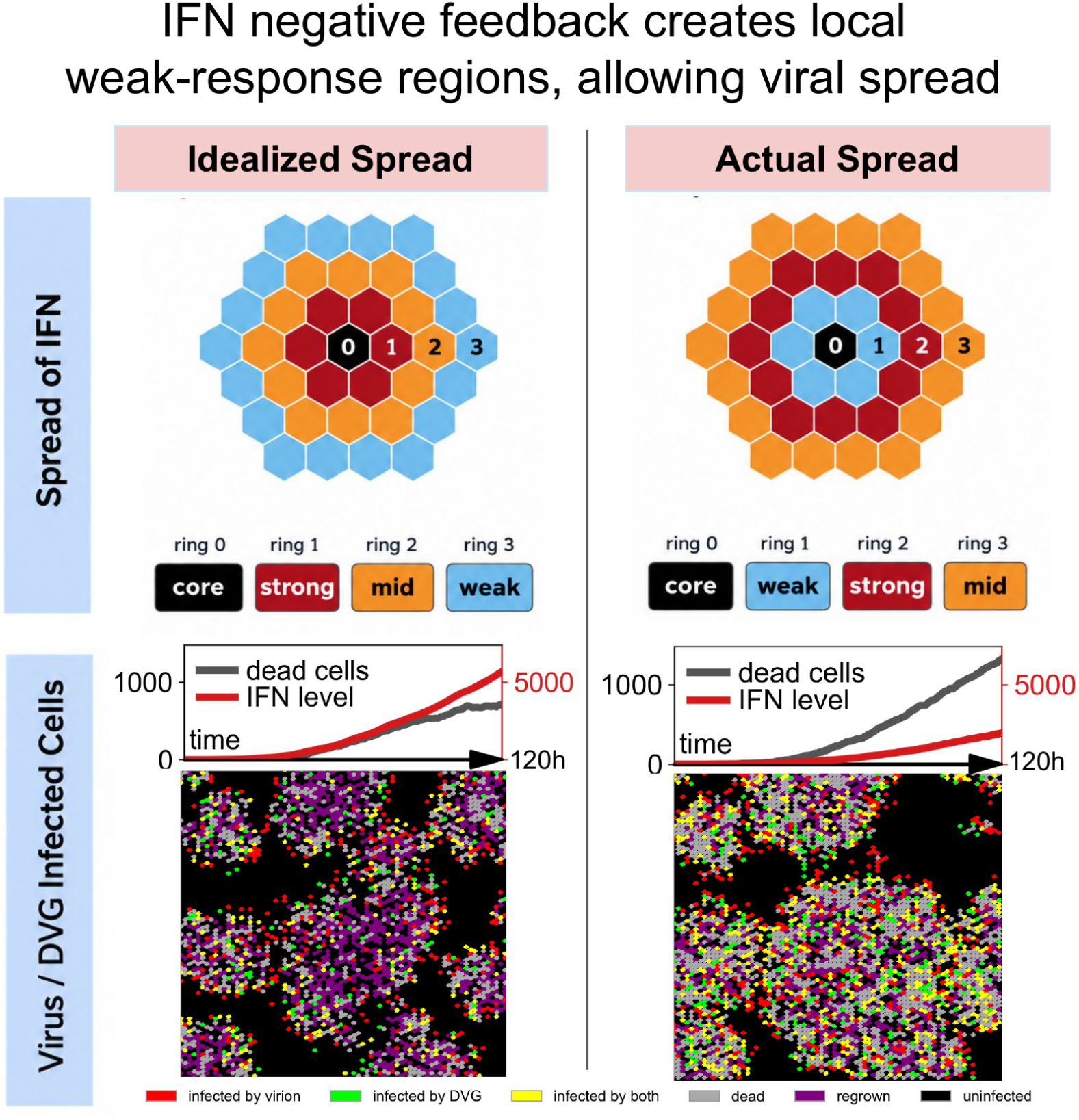
IFN negative feedback creates local weak-response regions, allowing viral spread. Top, spatial IFN response. Without negative feedback, IFN responses decrease with distance from the infection focus. With the non-monotonic response measured in our reporter experiment, cells nearest the focus show weaker subsequent IFN induction, while stronger responses occur farther away. **Bottom, spatial viral spread**. Stochastic simulations comparing monotonic and non-monotonic IFN regulation show more virus and DVG spread with negative feedback (one of the 30 runs per arm, shown as a representative, at 120 h; Methods, and the simulation movies on GitHub). The time course above each frame follows that same run: dead cells (grey, left axis, counts on the 76 *×*76 lattice) and IFN released summed over the run (red, right axis, model units). Spread on the lattice is between neighbours only; long-range jumps are not simulated, and how they would change this spatial pattern remains open.

To test this, we created a grid representing target cells in which the central cell represents the start of an infection. Without viral RNA-based stimulation of the negative feedback, we obtain a simple IFN gradient (Fig 5, top left). However, when we apply negative feedback by mapping our IFN dose-response curve onto the grid, the cells closest to the focus received the most secreted IFN but produced less subsequent IFN than cells located farther away (Fig 5, top right). Thus, while the extracellular IFN level may initially fall off from the site of infection as a simple gradient, the negative feedback modulates further IFN production in a spatial manner. This suppressed innate immune response in neighbouring cells could allow viruses to replicate and spread in a wave-like manner.

We further tested this spatial effect using stochastic simulations (Methods). The non-monotonic IFN response produced less IFN but greater viral spread (Fig 5, bottom), with the same pattern on a larger 360*×* 360 lattice (GitHub: `linear_video.mp4`, `nonlinear_video.mp4`, `nonlinear_VS_linear.png`).

This predicted ordering is consistent with measurements from viral microenvironments. Neighbouring cells can be more susceptible to secondary infection than more distal cells, despite their proximity to an infected cell. [61] However, whereas Song et al. attributed this pattern to factors transferred from infected cells, [61] our results suggest that IFN negative feedback alone may generate the same spatial pattern.

### Result 4: DVG abundance and IFN negative feedback shape responses to DVG-based and IFN-based interventions

DVG-based therapeutic strategies and IFN treatment have both been studied as antiviral interventions. [43, 44, 54, 62–65] However, DVG abundance can vary substantially between infections, [66] and the roles of virus-derived DVG, treatment-derived DVG, and IFN levels in shaping intervention outcome remain less well understood. We therefore used established forms of DVG and IFN intervention to perturb the modelled infection. Rather than identifying an optimal dose or timing, we asked how intervention effects change across different DVG and IFN regulatory conditions.

#### DVG intervention

Ideally, DVG treatment would suppress virus while increasing IFN (Fig 6, top). With cumulative-IFN feedback, however, the same intervention can produce different outcomes depending on the infection state at treatment (Fig 6, bottom). We therefore varied treatment time across no-, low-, and high-DVG conditions to probe how DVG–virus competition, IFN regulation, and target-cell availability interact (S7A Fig).

**Fig 6.**
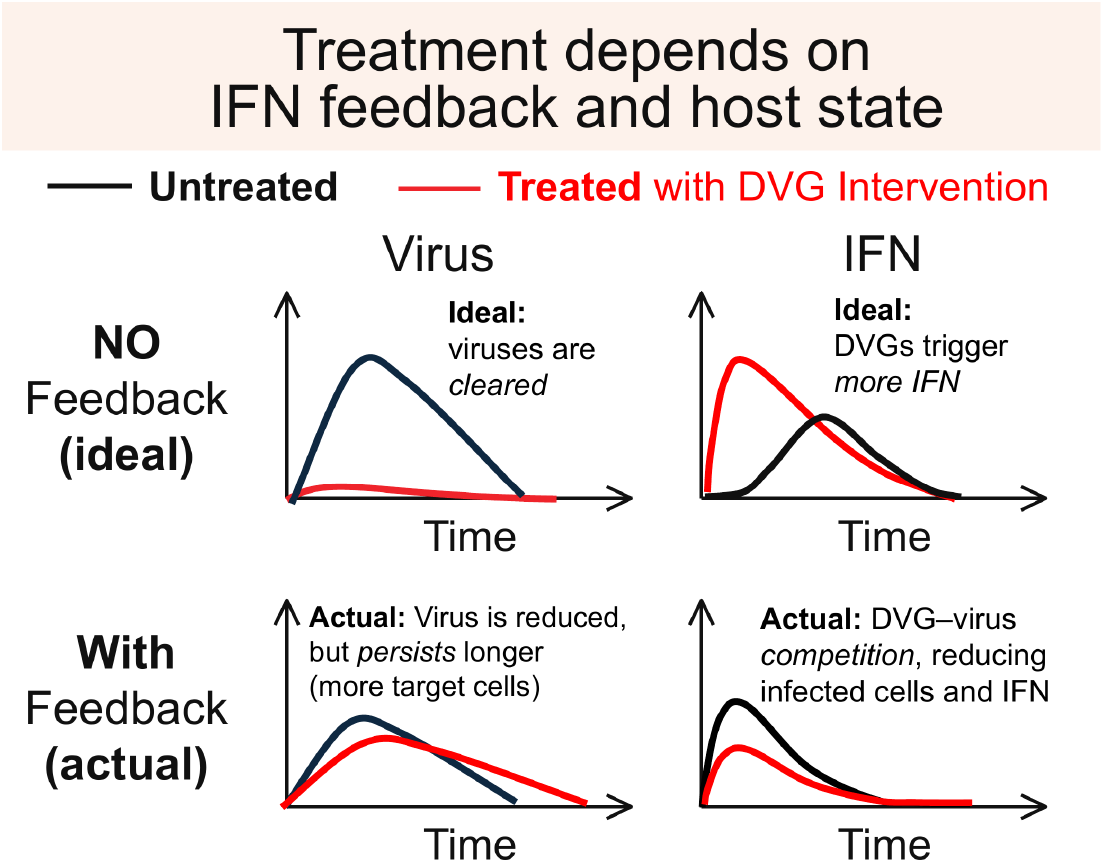
Schematic of DVG treatment with and without IFN negative feedback. Black, untreated; red, treated with DVGs. Top, the idealised outcome without feedback. Bottom, the outcome with cumulative-IFN feedback, as in our simulations. Curves are qualitative; simulated trajectories are in S7 Fig.

Early intervention is often expected to be most effective for DVG-based antivirals, which have frequently been studied as prophylactic or early treatments. However, our simulations exposed a trade-off between viral suppression and preservation of susceptible target cells. In the no- and low-DVG conditions, increasing DVG production before the viral peak shifted progeny away from virus and reduced early spread. This also preserved more susceptible cells, which could sustain infection later and extend the viral tail. Thus, treatment could lower the viral peak while increasing cumulative viral burden.

The IFN response depended on the pre-existing DVG state. In the no-DVG group, DVG-dependent IFN induction was not saturated, so treatment could produce a second IFN rise through stronger DVG sensing. In the low-DVG group, *s*(*D*) was already near saturation, so additional DVGs had little effect on IFN. The high-DVG group represented a later IFN regulatory state: IFN peaked early, cumulative negative feedback was already strongly engaged, and DVG-dependent induction was near saturation. Additional DVGs therefore had little effect on IFN after the earliest treatment.

Later intervention produced a simpler response. Once the viral peak had passed, the effects through IFN regulation and target-cell preservation became smaller, and DVG treatment acted mainly through DVG–virus competition to reduce subsequent infectious-virus production. This pattern was most evident by day 3 across all three DVG conditions.

Thus, these simulations do not identify a single optimal dose or timing. Instead, they show why the same DVG intervention can produce different outcomes depending on pre-existing DVG abundance, IFN history, DVG-induced IFN saturation, and the remaining target-cell pool. Peak virus, cumulative viral burden, and cell loss therefore respond differently to the same intervention timing.

#### IFN-based intervention

Ideally, IFN treatment should suppress virus directly. We therefore applied the same time-dependent antiviral effect across the no-, low-, and high-DVG conditions to ask how the surrounding IFN regulatory state changes the outcome (S7B Fig). Following previous IFN treatment studies, [65, 67, 68] we used a gradual reduction in virion production (maximum efficacy = 81%; 6-h half-rise; see Methods). [69, 70]

Early treatment again exposed a trade-off. In the no- and low-DVG groups, where cumulative IFN negative feedback remained weak, treatment reduced early virus but also preserved susceptible target cells. These cells could support infection later, extending the viral trajectory despite the lower early burden.

The high-DVG group was already in a different IFN regulatory state. Cumulative IFN exposure had strongly engaged negative feedback, so endogenous IFN production was already being suppressed. Treatment after this transition changed the IFN trajectory little and acted mainly by reducing subsequent virus production.

Hence, the same IFN intervention produced different outcomes depending on the pre-existing IFN state and remaining target-cell pool. Thus, treatment effects reflect not only antiviral efficacy itself, but also when the intervention enters the infection dynamics. The later treatment windows run long enough that an adaptive response would begin to arrive in a real host [71], which our innate-phase model does not carry; the late-treatment outcomes should be read with that in mind.

## Discussion

Our results use DVGs as a probe to understand how positive and negative regulation shape the type-I IFN response. We discuss how prior IFN exposure can suppress further IFN induction, and how this feedback shapes the timing and duration of the response. Finally, we consider its implications for DVG- and IFN-based interventions.

In basic models of the IFN response to viral infection [34, 40], IFN and virus both come from infected cells, so the two usually rise and fall together (Fig 2B, left). We here show that DVGs can break this coupling (Fig 3A and Fig 2E-H) and we built a whole-host IFN-feedback model that captures IFN induction (IFN-on) and cumulative IFN-based negative feedback (IFN-off). Our model and interpretations are supported by experiments (Fig 2E; [20, 37, 39]).

Beyond IAV, DVGs or DIPs have been linked to altered disease kinetics, antiviral or antitumour effects, and reduced performance of some live attenuated vaccines. [16, 39, 72, 73] Examples include measles virus, [74] rubella virus, [75] dengue virus, [76] other flaviviruses, [72] severe acute respiratory syndrome coronavirus 2 (SARS-CoV-2), [77, 78] respiratory syncytial virus, [23, 49] and other viruses. [73] DVGs also affect host cell-fate decisions across scales, from *in vitro* inference to within-host kinetics and population-level transmission models. [16, 79] DVG content therefore gives a broadly applicable probe for the IFN feedback *in vivo*.

Previous studies showed that an early IFN response can be protective, [20] while Rüdiger *et al*. showed how DVG interference can suppress infectious virus despite abundant viral RNA. [80] Our results add a third component: IFN induction itself can be transient even while the DVG-driven stimulus persists. Importantly, declining IFN concentration need not imply that the antiviral state established by the earlier response disappears on the same timescale. This distinction is consistent with the high-DVG condition, in which IFN declined earliest while virus remained low and mouse body-weight trajectories, the severity readout reported by Penn *et al*., resembled mock-infected controls, whereas the medium-DVG group sustained a larger IFN response and had poorer outcomes. [20] Thus, early IFN shutdown may limit prolonged inflammatory exposure without immediately removing the antiviral consequences established by the earlier response. Explicitly separating IFN concentration from a longer-lived antiviral state would provide a direct test of this hypothesis in future model extensions.

The cumulative-IFN threshold (cum*F*_50_) should be interpreted as an effective scale for IFN shutoff rather than as a single molecular threshold. It may combine molecular negative feedback, changes in IFN-producing cell states, and other unmeasured processes. Its identifiability also depends on whether the observed trajectory enters the feedback-sensitive range: the high-DVG data constrain this feedback most strongly, whereas the low- and medium-DVG trajectories provide substantially less information about the threshold.

We also refitted two classic influenza datasets that were first explained by host-side mechanisms alone: the equine pony data of Pawelek *et al*. [34] and the human titre data of Baccam *et al*. [40]. Virus–DIP competition reproduces both shapes, so DVG content is a candidate mechanism there too, alongside an IFN-dependent refractory return rate [46, 81] that acts on target-cell availability rather than on IFN induction (S3 Fig, S4 Fig).

In our simulations, the response to IFN treatment differed across the three DVG conditions, reflecting differences in the viral and IFN trajectories present when treatment began. This result does not establish DVG content as a determinant of IFN treatment response, but suggests that the infection state in which IFN is administered can matter. Defective HCV genomes have also been associated with relapse following pegylated IFN-*α* and ribavirin treatment, [82] although this clinical association does not establish either causality or the mechanism represented in our model. Testing whether DVG abundance contributes directly to IFN treatment response will require experiments in which DVG content and IFN exposure are independently controlled. Read together, these trade-offs are nonlinear in dose. In our simulations neither a large DVG dose nor a large IFN dose is reliably the best choice, and intermediate levels, judged against the host’s pre-existing IFN and DVG state, do better.

In our modelling, we focused on DIP datasets, in which DelVGs are likely the main drivers of innate immune activation. However, the feedback responds to accumulated IFN, and other noncanonical viral RNAs, such as mvRNAs produced by pathogenic IAV strains [13], but also agonists produced by bacterial co-infection [83] or bacteriophages [84], as well as IFN activation through mutations in viral antagonists [85] and to genetic host defects that impact the IFN response [86]. IFN gene expression also varies by cell type and by infection location along the respiratory tract [87, 88], so a change in DVG content should be read through its effect on the whole IFN trajectory, not by stronger immune activation alone. Previous studies have primarily characterised negative regulation at the receiving end of type-I IFN signalling, including regulation by USP18, SOCS proteins, and IFNAR. [2, 3, 32] Our reporter instead measures the production end: whether prior IFN exposure reduces subsequent IFN induction. By separating IFN exposure from infection, we show that prior IFN alone is sufficient to suppress further induction. The whole-host analysis further suggests that the history of IFN exposure, rather than its instantaneous level alone, can account for this regulation during infection.

### Caveats

Our model uses the cumulative-IFN term (*g*(cum*F* )) as a simple representation of how prior IFN exposure suppresses further IFN induction, rather than as a specific molecular mechanism such as USP18, SOCS proteins, or receptor internalisation. Experimentally observed IFN refractoriness can persist for roughly 72 h, [3, 89] making cumulative IFN exposure a reasonable approximation over our four-day data window. Over longer timescales, however, cells may recover from this refractory state, so cumulative exposure from the start of infection may no longer be an appropriate representation of IFN history. More generally, *g*(cum*F* ) is the simplest case of a memory kernel over past IFN exposure, one that weights early and recent exposure equally. Four timepoints over four days cannot separate that flat weighting from a kernel that discounts older exposure, so what is being stored up is not identified by these data.

Our reporter supports the central assumption behind this formulation: adding exogenous IFN lowers subsequent RNA-induced IFN-*β* promoter activation (Fig 2D,E), showing that prior IFN exposure can suppress further IFN induction. However, the reporter uses a single step dose of IFN rather than the gradual accumulation that occurs *in vivo*, and it does not identify which molecular processes carry this feedback. Thus, the experiment supports IFN-history-dependent suppression, but not the specific cumulative form used in the model.

Our fits track type-I IFN (IFN-*α in vivo*, IFN-*β in vitro*). Type-I IFN is antiviral signalling. It is distinct from the pro-inflammatory cytokines that mark an influenza cytokine storm, such as tumour necrosis factor (TNF) and interleukin-6 (IL-6) [6, 8]. Type-I IFN signalling can nevertheless contribute upstream to inflammatory pathology and cytokine production during influenza infection. [30] The same study also reported lung TNF [20]. TNF rose only in the medium-DVG group (7:1 Tky/05 LOW) at 48 hours. It stayed near baseline in the low- and high-DVG groups and at every other timepoint. This single elevated point is consistent with a transient cytokine-storm-like response in one group. It is too sparse to fit, and type-I IFN alone may not capture it. A fuller account of DVG-associated inflammation may need TNF and other cytokines together with type-I IFN.

We chose the simplest structure to focus on how DVG-dependent induction and cumulative IFN shutoff are constrained by the measured *V, D*, and *F* trajectories. Our model assigns IFN production to infected cells, although monocytes, macrophages, and plasmacytoid dendritic cells can also produce type-I IFN; [40] changes in these populations could therefore contribute to IFN decline independently of virus or DVG abundance. In addition, IFN was measured by ELISA on lung homogenate, so changes in bound or sequestered IFN could contribute to the measured decline. Although the inocula were matched by infectious dose, their total viral RNA content may differ.

Our model allows group-specific initial virus and DVG levels, but cannot distinguish DVG effects from those of other viral RNA components. Eclipse-phase formulations that separate infected cells into two states, or formulations in which virus competes with DVGs for cell entry (a varying *β*), also fit the data well in our ODE comparison (GitHub). We also keep the DVG production fraction (*d*) constant as a time-averaged approximation; our *in vitro* model treats this fraction explicitly over time [45].

Denser time series would allow stronger comparison among alternative equation forms. Longitudinal sampling within the same animals and absolute DVG copy counts would further tighten IFN constraints and reduce scaling uncertainty in *D*. Li *et al*. [45] report an *in vitro* time-course experiment (12, 24, 36, 48 hours; two to three exogenous IFN doses; reduced RNA grid) that tests the predicted pre-24 hours IFN peak and subsequent decline. We derive the reported *D* signal from DVG fractions and viral RNA and map it to a model state using a fixed scaling factor. The M-gene RNA measurements include both intracellular viral RNA and extracellular viral RNA in virions. Their relative contributions are unknown and may change over time. A larger intracellular share is expected early, while replication is active and released virions have yet to accumulate. Our *V* therefore tracks full-length viral RNA abundance rather than the released, infectious fraction a plaque assay reports. Fits to the Fig. 5e plaque-titre data are available on GitHub. Advances in quantification and consistent normalisation techniques across laboratories would improve inference and model comparison. [66, 90]

## Materials and Methods

### Study data

#### In vitro reporter assay

As shown in Fig 2C–E, we measured IFN-*β* promoter activation in HEK293-Luc cells, which carry a firefly-luciferase gene downstream of the IFN-*β* promoter. We use HEK293-Luc as a controlled IFN-*β* reporter to set IFN exposure independently of infection; the infection kinetics are measured in A549 lung epithelial cells. We transfected a triphosphorylated, double-stranded RIG-I ligand, as described previously [47, 48], and added exogenous type-I IFN at assay start. We varied the amount of transfected RIG-I agonist to give a graded IFN stimulus. This assay does not separate a direct effect on IFN-*β* promoter induction from an indirect effect through negative feedback. In our model, the turn-down of further induction follows from accumulated IFN rather than from the stimulus itself, and any RIG-I ligand that raises IFN can probe the same feedback. Luciferase was read 24 hours post-transfection as the endpoint; it reflects IFN-*β* promoter activation accumulated over the window, not the instantaneous IFN level. Repeats 1, 2, and 4 used RNA {0, 4, 20, 100, 500, 2500} ng crossed with exogenous IFN {0, 250, 500, 1000 ng/mL}, titrated by mass; the exogenous-IFN titration is shown in Fig 2F. Fig 2F shows the RNA ≤500 ng portion, each exogenous-IFN level normalised to its RNA = 0 baseline (fold change, baseline = 1) and combined across the three repeats by geometric mean; the RNA *>* 500 ng conditions enter a high-RNA regime analysed in separate work. We did not fit a dynamical model to this reporter assay.

#### In vivo mouse data

The BALB/c mouse experiment of Penn *et al*. [20] is, to our knowledge, currently unique for influenza A: a single *in vivo* study with time-resolved viral RNA, DVG level, and type-I IFN across inocula of different initial DVG content (Fig 3A). Adult female BALB/c mice (6–8 weeks) were infected intranasally with 10^5^ PFU of reassortant influenza A viruses carrying the six internal genes of A/turkey/Turkey/1/2005 (H5N1; Tky/05) and the hemagglutinin (HA) of A/Puerto Rico/8/34 (H1N1; PR8). The 6:2 Tky/05 (low-DVG) stock also carries the PR8 neuraminidase (NA), whereas the 7:1 Tky/05 LOW and HIGH stocks retain the Tky/05 NA. This reassortant background was used because the HPAIV H5N1 internal genes drive cytokine-mediated immunopathology more faithfully than well-characterised laboratory strains such as PR8; the H5N1 NS segment encodes the type-I IFN antagonist NS1, though IFN evasion of these stocks was not separately characterised [20]. The stocks were propagated at different multiplicity of infection (MOI) and differ in genome-to-PFU ratio, which sets the inoculum DVG content. We read 6:2 Tky/05, 7:1 Tky/05 LOW, and 7:1 Tky/05 HIGH as low, intermediate, and high initial DVG; the low-DVG stock was passaged in Madin-Darby canine kidney (MDCK) cells at MOI 0.0001 to minimise DVG generation. This design lets us test how the starting DVG level affects the joint kinetics of virus and IFN, and estimate the DVG contribution by comparing no-DVG and with-DVG fits. Five mice per group were sampled at each time point; lungs were harvested and homogenised, and infectious virus (plaque assay), total viral RNA (M-gene RT-qPCR), and IFN-*α* (mouse IFN-*α* ELISA; VeriKine, PBL) were measured in the lung homogenate [20].

The IFN-*α* kit (VeriKine, PBL, catalogue 42120) is specified by its manufacturer over an assay range of 12.5–400 pg mL^*−*1^, with the lowest standard of its six-point curve at 12.5 pg mL^*−*1^ [91]; Penn *et al*. report no detection limit for this measurement.

#### Data processing

Each group provides virus *V*, DVG load *D*_obs_ and IFN-*α F* (pg mL^*−*1^) at 6, 24, 48 and 96 hours, five mice at each. Virus and DVG are both counted as genome copies per gram, full-length and defective respectively, so the two are in one unit. Both are read from the per-animal M-gene measurements of Penn *et al*. Fig 5f, not from the plaque titres of their Fig 5e, and *F* is read from their Fig 6b. *V* and *D*_obs_ enter as log_10_; *F* enters on the linear pg mL^*−*1^ scale it is measured on. For *D*_obs_ we use the DVG fractions in the polymerase segments (PB1, PB2, PA), which carry 97.6% of the junction reads in this experiment and are where DVGs are most often detected [10, 92]: we average the three junction-read fractions, taken from

Table S1 of their supplemental material, multiply by total viral RNA (M-gene copies) to obtain an amount proportional to absolute DVG copies, and take log_10_. Reading that fraction as the fraction of genomes that are defective is an assumption; Penn *et al*. note that the read-based measure supports relative comparisons between samples rather than accurate DVG frequencies. This estimated DVG level is linked to the model state *D* by a scaling constant *m*_1_ (Supporting Information): DVGs are genomes, DIPs are the particles those genomes are packaged into. The with-DVG model is fit to *V, D*_obs_, and *F* ; the no-DVG model to *V* and *F* only. The measured species is IFN-*α in vivo* and IFN-*β in vitro*^1^, and the Results use the generic “IFN” (Fig 3A).

#### In vitro A549 data

The same study also reports A549 *in vitro* time courses of standard viral genomes, DVG genomes, and IFN-*β* mRNA for the same three stocks, sampled at 2, 6, and 24 hours. A549 are human alveolar (lung) epithelial cells [20]. We fit the same with-DVG and no-DVG models to these by the same procedure. The fits show a compensation pattern *in vitro*: without an explicit DVG state, the no-DVG model raises the infected-cell loss rate *δ* to match the data, most strongly in the high-DVG condition (Supporting Information, S2 and S5 Figs).

### Model and fitting

We fit two model structures to each data set (Fig 3B): a with-DVG structure (*T, I, V, D, F* ) and a no-DVG structure that drops the DVG state together with the parameters *d, h, D*_0_, *s*_*d*_, and *θ*. The with-DVG structure is fit with and without cumulative-IFN feedback, and the no-DVG structure with it, giving three variants. Cumulative IFN exposure is cum 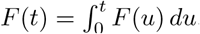. The feedback term *g*(cum*F* ) = 1*/*(1 + cum*F/*cum*F*_50_) lets accumulated IFN reduce further induction, matching how IFN signalling limits itself: IFN induces negative regulators such as SOCS proteins and USP18, and reduces signalling through IFNAR downregulation and internalisation. [2] We chose the form to reproduce the measured timing of the virus and IFN trajectories, not to assert that one molecular species was measured directly. We use “IFN” for type-I IFN throughout. The main with-DVG model with cumulative-IFN feedback is defined in Results (Eqs. 1–3 and Table 1); the variants and the fitting procedure are described here.

#### Base model with both positive and negative IFN feedback

Removing the DVG compartment from Eq. 1, by setting *d* = 0 and *D*(0) = 0 so that *s*(*D*) reduces to *s*_0_, gives the core whole-host model with both IFN feedbacks and no DVG pathway, drawn as the inner *Model with IFN feedback* box of Fig 3B:

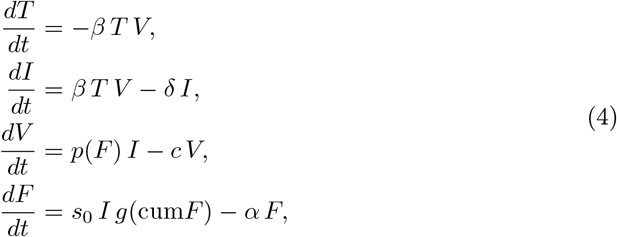

with initial conditions *T* (0) = *T*_0_, *I*(0) = 0, *V* (0) = *V*_0_, *F* (0) = 0, cum*F* (0) = 0, and with 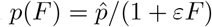 and *g*(cum*F* ) as in the full model. This variant is used for the no-DVG fits below and in the Supporting Information. Dropping *F* in turn leaves the classical target-cell-limited backbone (innermost box of Fig 3B [40]).

#### With-DVG fit without cumulative-IFN feedback

Removing feedback drops the cum*F* state and sets *g* ≡ 1. The states are *T, I, V, D*, and *F* :

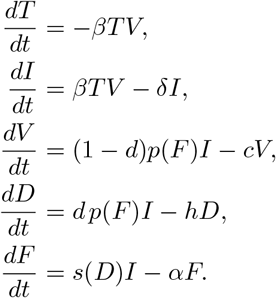

#### No-DVG fit with cumulative-IFN feedback

Removing DVGs drops the *D* state and the parameters *d, h, D*_0_, *s*_*d*_, and *θ*; induction collapses to *s*_0_ and *V* loses the (1 −*d*) factor. This is the base model of Eq. 4 (Fig 3B, *Model with IFN feedback* box), written here with the cumulative-exposure state explicit. The states are *T, I, V, F*, and cum*F* :

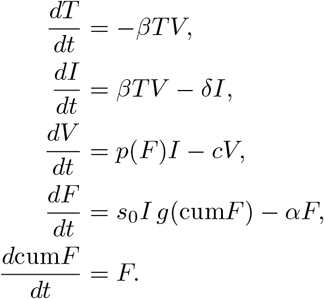

#### Testing the IFN feedback term

We evaluated the contribution of this feedback by comparing the full model with a no-feedback model (*g*(cum*F* ) ≡ 1). We also profiled cum*F*_50_ separately for the low-, medium-, and high-DVG groups to determine how strongly each dataset supported the feedback term. Model parameters were re-estimated for each comparison.

#### Alternatives

Alternative *s*(*D*) forms and ODE structures were fit to the same data in the project repository (GitHub). In particular, a refractory-cell recovery path that returns IFN-protected cells to the susceptible pool [34] can be written as an alternative equation, but it does not reproduce the high-DVG IFN decline either.

Refractory recovery refills the susceptible pool *T*, which permits further infection and keeps the infected-cell pool *I* from collapsing at exactly the time *V* and *D* are still rising. With *s*(*D*) *I* as the IFN source, this sustains IFN production rather than shutting it off. *F* therefore does not fall to near-zero as observed in the high-DVG group at 24–48 hours. The cumulative form is preferred by fit; a current-level feedback *g*(*F* ) does not reproduce the high-DVG IFN decline even at unreasonably high *α*.

#### Inference

The three groups differ in their inocula, so each model is fit jointly across them: host-related parameters are shared across groups, whereas the remaining parameters are fit separately for each group (fits with all parameters estimated separately by group are also available on GitHub). Inference is in Stan [93] with the No-U-Turn Sampler (NUTS; 4 chains, 2,000 warm-up and 4,000 sampling iterations); below-detection values are left-censored. The parameterisation enforces *c > δ* and (with-DVG) *h > δ*, so virion and DVG clearance always exceed infected-cell loss. The prior on the type-I IFN decay rate (*α*) is informed by published estimates from within-host influenza models. [33, 34] A weakly-informative log-normal prior on 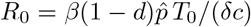 (the no-DVG variant drops the (1 −*d*) factor) constrains the joint identifiability of 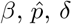, and *c*. Posterior medians with 90% credible intervals are reported in S1 Table for the with-DVG model, and on GitHub for the no-DVG model, and the prior families and bounds in S2 Table. Initial conditions are *T*_0_, *V*_0_, *D*_0_ (with-DVG only), *I*_0_ = *F*_0_ = 10^*−*10^ (numerical zero), and cum*F* (0) = 0 for the feedback variants. DVG measurements provide additional information but are not required to apply the cumulative-IFN feedback framework. DVG abundance is not usually represented by standard measures of infectious virus and requires additional quantification. [16] Importantly, when DVG measurements are unavailable, the no-DVG model fitted to virus and IFN alone recovers the same qualitative cumulative-feedback pattern as the DVG-resolved model. Thus, DVG measurements add mechanistic resolution, but the feedback framework does not depend on them.

A simulation-based recovery check refits the model to synthetic data generated at the fitted posterior medians. The model recovers 38 of the 39 estimated quantities within their 90% credible intervals; the remaining quantity is also closely recovered (S6 Fig).

#### Spatial simulation

We used the experimentally calibrated stochastic spatial model of influenza A virus, DVGs, and type-I IFN developed by Li *et al*. [45] (code and run scripts on GitHub). The spatial structure, infection process, IFN spread, and associated parameters follow the original model.

We compared two simulation arms that differ only in how much IFN a cell produces for the viral RNA it carries. The first uses the constant IFN output of the original model. The second uses a non-monotonic IFN response scaled to the relative response measured in our reporter experiments. All other model assumptions and parameters were kept unchanged (details in Li *et al*. [45] and GitHub).

We compared the two arms across independent stochastic runs (30 seeds per arm, 76× 76 lattice, 20 initial infection foci). Fig 5 shows the median realisation from each arm at 120 h. Larger visualization runs used a 360× 360 lattice with 20 initial foci (movies on GitHub).

#### DVG and IFN intervention simulations

We simulated IFN- and DVG-based interventions beginning on days 0, 1, 3, or 8 over a 15-day period, on the fitted model with no additional terms. The no-DVG condition is the low-DVG group’s no-DVG posterior with *d* = 0 and *D*(0) = 0, carrying *h, s*_*d*_ and *θ* from randomly paired draws of that group’s with-DVG run.

The IFN intervention takes the antiviral-efficacy form used in the hepatitis C literature: total progeny production carries a time-dependent factor 1 −*e*(*t*) with *e*(*t*) = 0.81 {1 −exp[− *k*(*t*− *τ* )] } for *t* ≥ *τ* and *k* = ln(2)*/*(6*/*24) day^*−*1^. The maximum efficacy of 81% is the reported estimate for interferon alfa-2b treatment, [65] and the gradual onset follows an established time-varying efficacy form, [69, 70] for which we set a 6-h half-rise. The factor applies equally to infectious-virus and DVG production. No interferon is added to *F*, so the treatment leaves both endogenous IFN and cumulative IFN exposure untouched, and it continues for the remainder of the simulation.

The DVG intervention adds a single dose (100 times the fitted high-DVG inoculum) at the intervention time and holds the fraction of progeny produced as DVGs at *d*_treat_ until day 15. We set that fraction per group to *d*_treat_ = 1− (1− 0.81)(1− *d*_50_), where *d*_50_ is the median of *d* across the 200 draws, which gives 0.810, 0.810 and 0.902 for the no-DVG, low-DVG and high-DVG conditions. At that median draw the two interventions leave the same infectious-virus production. What separates them is the rest of the output: the DVG intervention leaves total progeny production unchanged and sends that remainder to the defective compartment, where it is cleared at rate *h* and feeds IFN induction through *s*(*D*), while the IFN intervention removes it. Defective production is therefore not matched, the DVG intervention sending a share 0.810, 0.810 and 0.902 of production to *D* against 0, 0.0001 and 0.092 under the IFN intervention. The match on virus also holds only as closely as *d* is determined within a group: exactly in the no-DVG condition, where *d* = 0 in every draw, and to within 1% across the central 90% of low-DVG draws, but the high-DVG posterior for *d* runs from 0.04 to 0.97 over that same range, and in 14% of its draws the fitted *d* already exceeds *d*_treat_, so the intervention lowers their defective fraction. These values are large enough to separate the treated trajectory from the untreated one on a log_10_ axis, and they stay below the near-complete defective-progeny production reported for co-infected cells [53].

Trajectories were simulated across 200 posterior draws and summarised at each time point by averaging the middle 80% of trajectories after removing the highest and lowest 10%.

## Declarations

### Authors’ contributions

Y.L.: conceptualization, methodology, software, validation, formal analysis, writing – original draft, visualization, writing – review and editing; C.L.: methodology, validation, investigation; B.F.N.: validation, software, formal analysis, writing – review and editing; S.A.L.: writing – review and editing, funding acquisition; A.S.P.: validation, formal analysis, writing – review and editing; A.J.W.t.V.: methodology, validation, formal analysis, supervision, writing – review and editing, funding acquisition; B.T.G.: formal analysis, funding acquisition, supervision, validation, writing – review and editing.

All authors gave final approval for publication and agreed to be held accountable for the work performed therein.

## Acknowledgements

We thank Alice S. Huang for discussions on defective interfering particles. We thank Sarah Cobey for helpful comments on the manuscript. We thank Hwai-Ray Tung and Emily Howerton for helpful discussions.

## Data and code availability

All data supporting the results of this study are openly available at: https://github.com/yimei-li/within-host-dip-iav-ode.

## Ethics statement

This study did not involve human participants. The *in vitro* reporter experiments used an immortalised cell line (HEK293-Luc) and did not require ethical approval. The mouse and A549 data are a secondary analysis of previously published experiments [20], for which ethical approval was obtained in the original study. No new animal experiments were performed for this work.

## Use of Artificial Intelligence (AI)-assisted technologies

AI-assistance was used for grammar correction and completing lines in our code. All AI-produced code and information were verified by Yimei Li.

## Supporting Information

**Table S1.**
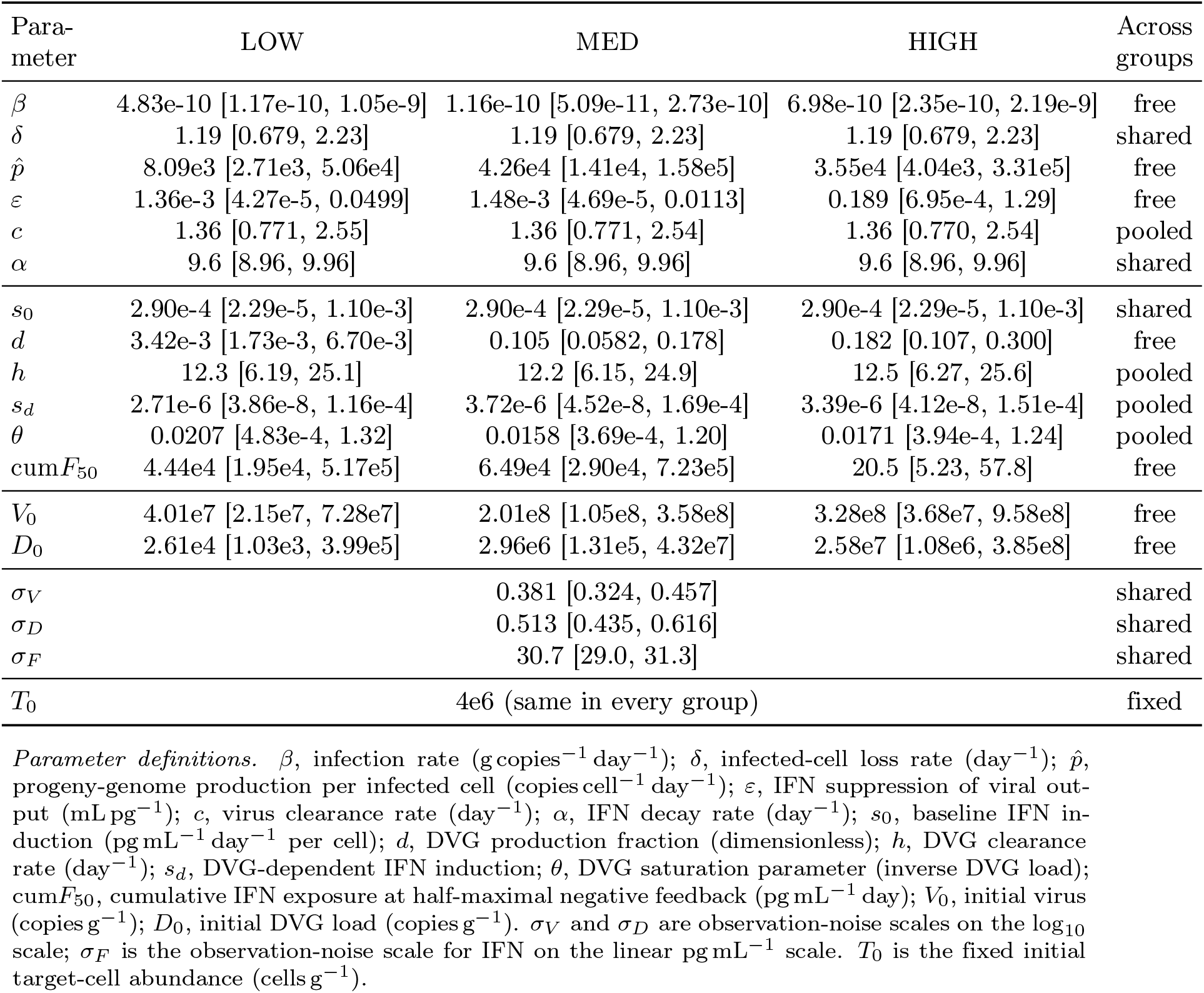
Posterior medians with 90% credible intervals [5%, 95%] for the with-DVG model across inoculum groups. The three groups are fitted jointly: host-related parameters are shared or partially pooled across groups, whereas inoculum- and DVG-related parameters are estimated separately by group, as indicated in the final column. IFN dynamics include induction and cumulative-IFN negative feedback; below-LOD observations are left-censored. Priors for the infected-cell loss rate (*δ*), virus clearance rate (*c*), and IFN decay rate (*α*) were informed by published within-host influenza estimates [33, 34, 40, 71]; all prior distributions and bounds are given in Table S2. In the low- and medium-DVG groups, cumulative IFN exposure remains below the range in which negative feedback is appreciably engaged; accordingly, the data primarily constrain *g*(cum*F* ) to remain near one rather than precisely identifying cum*F*_50_. The large cum*F*_50_ estimates in these groups therefore represent weak feedback over the observed time window rather than precisely estimated thresholds. In the high-DVG group, cumulative IFN exposure enters the feedback-sensitive range, allowing cum*F*_50_ to be more directly constrained.

| Para-meter | LOW | MED | HIGH | Across groups |
|---|---|---|---|---|
| β | 4.83e-10 [1.17e-10, 1.05e-9] | 1.16e-10 [5.09e-11, 2.73e-10] | 6.98e-10 [2.35e-10, 2.19e-9] | free |
| δ | 1.19 [0.679, 2.23] | 1.19 [0.679, 2.23] | 1.19 [0.679, 2.23] | shared |
| *p̂* | 8.09e3 [2.71e3, 5.06e4] | 4.26e4 [1.41e4, 1.58e5] | 3.55e4 [4.04e3, 3.31e5] | free |
| ε | 1.36e-3 [4.27e-5, 0.0499] | 1.48e-3 [4.69e-5, 0.0113] | 0.189 [6.95e-4, 1.29] | free |
| *c* | 1.36 [0.771, 2.55] | 1.36 [0.771, 2.54] | 1.36 [0.770, 2.54] | pooled |
| α | 9.6 [8.96, 9.96] | 9.6 [8.96, 9.96] | 9.6 [8.96, 9.96] | shared |
| *s*<sub>0</sub> | 2.90e-4 [2.29e-5, 1.10e-3] | 2.90e-4 [2.29e-5, 1.10e-3] | 2.90e-4 [2.29e-5, 1.10e-3] | shared |
| *d* | 3.42e-3 [1.73e-3, 6.70e-3] | 0.105 [0.0582, 0.178] | 0.182 [0.107, 0.300] | free |
| *h* | 12.3 [6.19, 25.1] | 12.2 [6.15, 24.9] | 12.5 [6.27, 25.6] | pooled |
| *s*<sub>*d*</sub> | 2.71e-6 [3.86e-8, 1.16e-4] | 3.72e-6 [4.52e-8, 1.69e-4] | 3.39e-6 [4.12e-8, 1.51e-4] | pooled |
| θ | 0.0207 [4.83e-4, 1.32] | 0.0158 [3.69e-4, 1.20] | 0.0171 [3.94e-4, 1.24] | pooled |
| cum*F*<sub>50</sub> | 4.44e4 [1.95e4, 5.17e5] | 6.49e4 [2.90e4, 7.23e5] | 20.5 [5.23, 57.8] | free |
| *V*<sub>0</sub> | 4.01e7 [2.15e7, 7.28e7] | 2.01e8 [1.05e8, 3.58e8] | 3.28e8 [3.68e7, 9.58e8] | free |
| *D*<sub>0</sub> | 2.61e4 [1.03e3, 3.99e5] | 2.96e6 [1.31e5, 4.32e7] | 2.58e7 [1.08e6, 3.85e8] | free |
| σ<sub>*V*</sub> | 0.381 [0.324, 0.457] |  |  | shared |
| σ<sub>*D*</sub> | 0.513 [0.435, 0.616] |  |  | shared |
| σ<sub>*F*</sub> | 30.7 [29.0, 31.3] |  |  | shared |
| *T*<sub>0</sub> | 4e6 (same in every group) |  |  | fixed |

*Parameter definitions.* β, infection rate (g copies<sup>−1</sup> day<sup>−1</sup>); δ, infected-cell loss rate (day<sup>−1</sup>); *p̂*, progeny-genome production per infected cell (copies cell<sup>−1</sup> day<sup>−1</sup>); ε, IFN suppression of viral out-put (mL pg<sup>−1</sup>); *c*, virus clearance rate (day<sup>−1</sup>); α, IFN decay rate (day<sup>−1</sup>); *s*<sub>0</sub>, baseline IFN in-duction (pg mL<sup>−1</sup> day<sup>−1</sup> per cell); *d*, DVG production fraction (dimensionless); *h*, DVG clearance rate (day<sup>−1</sup>); *s*<sub>*d*</sub>, DVG-dependent IFN induction; θ, DVG saturation parameter (inverse DVG load); cum*F*<sub>50</sub>, cumulative IFN exposure at half-maximal negative feedback (pg mL<sup>−1</sup> day); *V*<sub>0</sub>, initial virus (copies g<sup>−1</sup>); *D*<sub>0</sub>, initial DVG load (copies g<sup>−1</sup>). σ<sub>*V*</sub> and σ<sub>*D*</sub> are observation-noise scales on the log<sub>10</sub> scale; σ<sub>*F*</sub> is the observation-noise scale for IFN on the linear pg mL<sup>−1</sup> scale. *T*<sub>0</sub> is the fixed initial target-cell abundance (cells g<sup>−1</sup>).

### Inference details: priors, likelihood, initial conditions, and credible intervals

This section records the prior families, the observation likelihood, the fixed initial conditions, and the credible-interval definition, so the fits can be reproduced from the deposited data alone. Numeric values are the *in vivo* settings; the *in vitro* fits use the same prior families and likelihood with data-scaled hyperparameters.

#### Priors

Each kinetic rate has a log-normal prior, written log(·) ∼ Normal(*µ, σ*) on the log parameter and truncated to the bounds in S2 Table. The DVG fraction uses a logit-normal prior, logit(*d*) ∼ Normal(*µ*_*d*_, 3), and the initial DVG load *D*_0_ a log-normal prior. The observation-noise scales *σ*_*V*_ and *σ*_*D*_ use Exponential(0.5), and *σ*_*F*_ uses Exponential(0.0319). A weakly-informative log-normal prior on 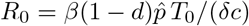 (the no-DVG variant drops the 1 *d* factor), centred at 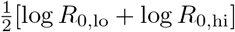 with *R*_0,lo_ = 1, *R*_0,hi_ = 200 and scale 2, links 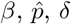, and *c*. The parameterisation enforces *c > δ* and *h > δ*.

#### Likelihood

Virus and DVG enter on the log_10_ scale, IFN on the linear pg mL^*−*1^ scale it is measured on. Virus and DVG use a normal likelihood, *y*_*V*_∼ Normal(log_10_ *V, σ*_*V*_ ) and *y*_*D*_∼ Normal(log_10_(*D/m*_1_), *σ*_*D*_), where *y*_*D*_ is the estimated DVG level log_10_ *D*_obs_ and *m*_1_ links it to the model state as *D* = *m*_1_*D*_obs_. Here *m*_1_ = 1: virus is counted as full-length genome copies per gram and DVG as defective genome copies per gram, so the two are already in one unit and no conversion enters. The split of output by *d* and 1− *d* is then between commensurate quantities, and the values reported for *D, D*_0_ and *θ* are on that genome scale, in copies g^*−*1^. This differs from the earlier fits of this model, which measured virus as PFU g^*−*1^ and carried *m*_1_ = 5 ×10^*−*5^ to put the two on a common particle scale; the constant is retained in the likelihood so the two readings share one expression. IFN uses a normal likelihood on the linear scale, *y*_*F*_ Normal(*F, σ*_*F*_ ), with *σ*_*F*_ bounded above by 31.3 pg mL^*−*1^, the within-timepoint standard deviation of the measurements themselves. Below-detection observations are left-censored: the contribution is the lower-tail probability of the same distribution evaluated at the detection limit. The limits of detection (LOD) are LOD_*V*_ = LOD_*D*_ = 0 on the log_10_ scale, and LOD_*F*_ = 12.5 pg mL^*−*1^ on the linear one. Only IFN observations fall below their limit: 14 of 20 in low, 10 of 20 in med and 14 of 20 in high. The IFN limit is the lowest concentration the assay quantifies rather than a figure Penn *et al*. report: they state no detection limit for the lung-homogenate IFN-*α* ELISA, and the manufacturer specifies the assay over 12.5–400 pg mL^*−*1^ [91], the lowest point of its six-point standard curve being 12.5 pg mL^*−*1^. Censoring there says of an unquantifiable animal only what the instrument supports, that it lies below the working range. Earlier fits of this model censored at 10^*−*2^ pg mL^*−*1^, three orders below that point, which instead entered such an animal as though it were known to be near zero.

#### Initial conditions

The target-cell pool starts at *T*_0_ = 4 ×10^6^ *in vivo* and 1×10^6^ *in vitro*. The main text states the initial infected-cell and IFN states as *I*(0) = 0 and *F* (0) = 0. In the fits these are set to a value very close to zero, so that the solver and the log_10_ observation model stay defined at *t* = 0; infection is still seeded by *V*_0_ through the *βTV* term, so the floor carries no biological content. cum*F* (0) = 0. Initial virus *V*_0_ and (with-DVG) initial DVG load *D*_0_ are fitted (S1 Table).

#### Credible intervals

Shaded bands are pointwise posterior quantiles: the 50% band is [25%, 75%] and the 90% band is [5%, 95%]. Tabulated intervals are the [5%, 95%] quantiles. These are quantiles of the posterior, not normal approximations.

**Table S2.**
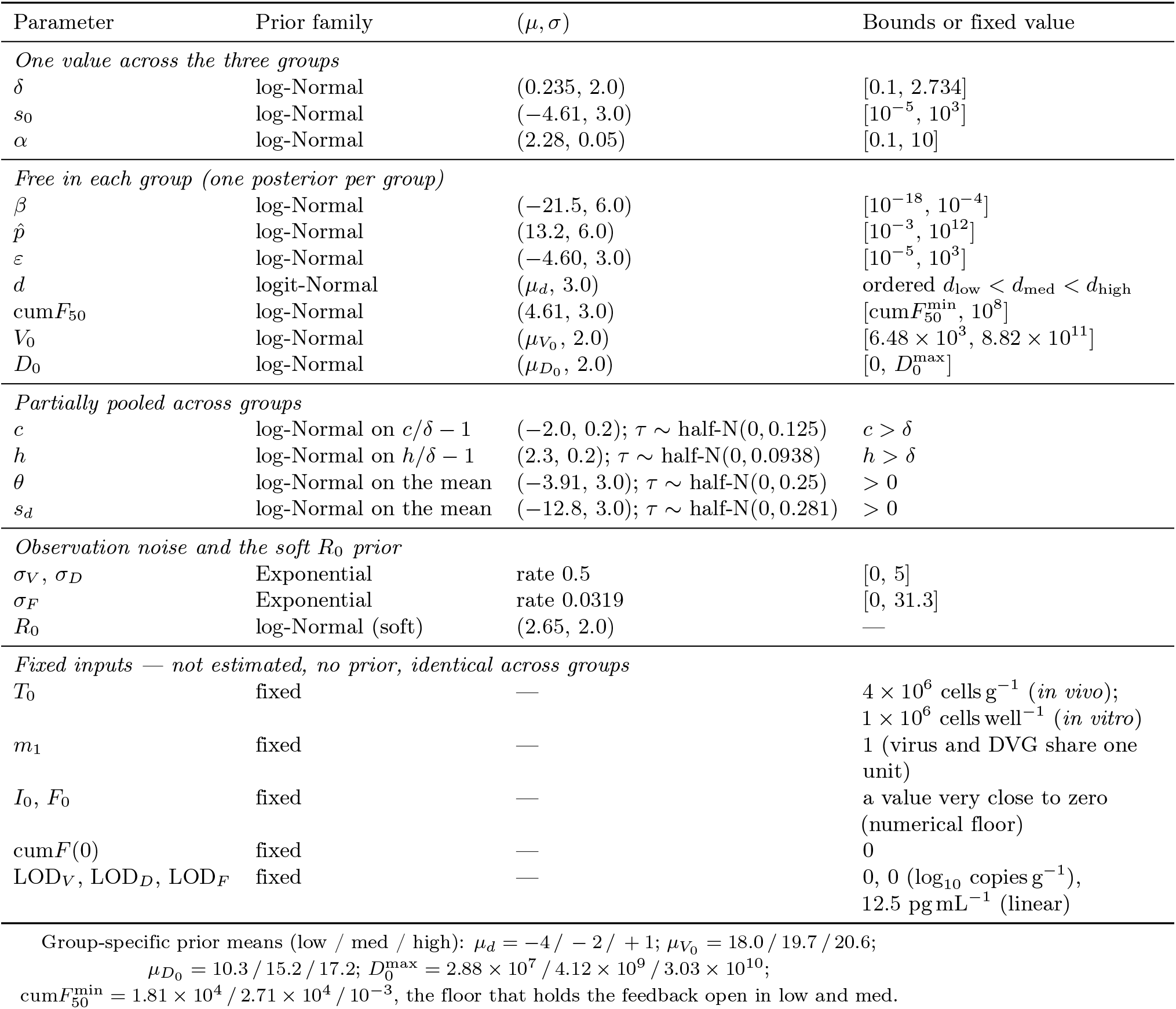
Prior distributions and bounds for the with-DVG model (*in vivo*). log-Normal (*µ, σ*) denotes a normal prior on the log parameter, truncated to the bounds shown; logit-Normal denotes a normal prior on logit *d*, which is unbounded. Bounds give the support of the truncated prior. The no-DVG model drops *d, h, s*_*d*_, *θ, D*_0_ and *σ*_*D*_. Group-specific prior means (low / med / high) are listed below the table where they differ; the *in vitro* fits share these families, though not every bound: the tabled numeric bounds are the *in vivo* settings, and the *in vitro* fits rescale some of them to their data (for example the upper bound on *β*). The blocks separate the four kinds of input. Parameters in the first block take one value across the three groups. Those in the second are free in each group, with one posterior per group. Those in the third are partially pooled: the group values are drawn around a common mean whose spread *τ* carries its own half-normal prior, so they are neither tied together nor independent. *c* and *h* carry their prior on the ratio to *δ* rather than on the rate itself, which is what makes the orderings *c > δ* and *h > δ* hold in every group with no rejection step. *R*_0_ is not a free parameter but a function of 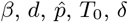 and *c* carrying a soft prior that links them. Rows in the last block are fixed before fitting, take no prior, and are held at the same value in every group, so no posterior is reported for them; *T*_0_ and *m*_1_ are structural choices rather than literature estimates, *I*_0_ and *F*_0_ are numerical floors (see Initial conditions), and the detection limits come from the assays. *m*_1_ = 1 here because virus and DVG are both counted as genome copies per gram, so no genome-to-particle conversion enters. Reasons for the two rate orderings are given under Priors.

| Parameter | Prior family | (μ, σ) | Bounds or fixed value |
|---|---|---|---|
| *One value across the three groups* |  |  |  |
| δ | log-Normal | (0.235, 2.0) | [0.1, 2.734] |
| s<sub>0</sub> | log-Normal | (−4.61, 3.0) | [10<sup>−5</sup>, 10<sup>3</sup>] |
| α | log-Normal | (2.28, 0.05) | [0.1, 10] |
| *Free in each group (one posterior per group)* |  |  |  |
| β | log-Normal | (−21.5, 6.0) | [10<sup>−18</sup>, 10<sup>−4</sup>] |
| p̂ | log-Normal | (13.2, 6.0) | [10<sup>−3</sup>, 10<sup>12</sup>] |
| ε | log-Normal | (−4.60, 3.0) | [10<sup>−5</sup>, 10<sup>3</sup>] |
| d | logit-Normal | (μ<sub>d</sub>, 3.0) | ordered d<sub>low</sub> < d<sub>med</sub> < d<sub>high</sub> |
| cumF<sub>50</sub> | log-Normal | (4.61, 3.0) | [cumF<sub>50</sub><sup>min</sup>, 10<sup>8</sup>] |
| V<sub>0</sub> | log-Normal | (μ<sub>V<sub>0</sub></sub>, 2.0) | [6.48 × 10<sup>3</sup>, 8.82 × 10<sup>11</sup>] |
| D<sub>0</sub> | log-Normal | (μ<sub>D<sub>0</sub></sub>, 2.0) | [0, D<sub>0</sub><sup>max</sup>] |
| *Partially pooled across groups* |  |  |  |
| c | log-Normal on c/δ − 1 | (−2.0, 0.2); τ ∼ half-N(0, 0.125) | c > δ |
| h | log-Normal on h/δ − 1 | (2.3, 0.2); τ ∼ half-N(0, 0.0938) | h > δ |
| θ | log-Normal on the mean | (−3.91, 3.0); τ ∼ half-N(0, 0.25) | > 0 |
| s<sub>d</sub> | log-Normal on the mean | (−12.8, 3.0); τ ∼ half-N(0, 0.281) | > 0 |
| *Observation noise and the soft R<sub>0</sub> prior* |  |  |  |
| σ<sub>V</sub>, σ<sub>D</sub> | Exponential | rate 0.5 | [0, 5] |
| σ<sub>F</sub> | Exponential | rate 0.0319 | [0, 31.3] |
| R<sub>0</sub> | log-Normal (soft) | (2.65, 2.0) | — |
| *Fixed inputs — not estimated, no prior, identical across groups* |  |  |  |
| T<sub>0</sub> | fixed | — | 4 × 10<sup>6</sup> cells g<sup>−1</sup> (*in vivo*); 1 × 10<sup>6</sup> cells well<sup>−1</sup> (*in vitro*) |
| m<sub>1</sub> | fixed | — | 1 (virus and DVG share one unit) |
| I<sub>0</sub>, F<sub>0</sub> | fixed | — | a value very close to zero (numerical floor) |
| cumF(0) | fixed | — | 0 |
| LOD<sub>V</sub>, LOD<sub>D</sub>, LOD<sub>F</sub> | fixed | — | 0, 0 (log<sub>10</sub> copies g<sup>−1</sup>), 12.5 pg mL<sup>−1</sup> (linear) |

Group-specific prior means (low / med / high): μ<sub>d</sub> = −4 / − 2 / + 1; μ<sub>V<sub>0</sub></sub> = 18.0 / 19.7 / 20.6; μ<sub>D<sub>0</sub></sub> = 10.3 / 15.2 / 17.2; D<sub>0</sub><sup>max</sup> = 2.88 × 10<sup>7</sup> / 4.12 × 10<sup>9</sup> / 3.03 × 10<sup>10</sup>; cumF<sub>50</sub><sup>min</sup> = 1.81 × 10<sup>4</sup> / 2.71 × 10<sup>4</sup> / 10<sup>−3</sup>, the floor that holds the feedback open in low and med.

#### Data availability

The processed virus, DVG, and IFN time series fed to every fit, the fixed model inputs (*T*_0_, *m*_1_, *F*_0_, detection limits), and the posterior-median tables for all fits are deposited as plain CSV files in the project repository (https://github.com/yimei-li/within-host-dip-iav-ode).

### Analytic argument: the IFN decline requires negative feedback

The model with positive IFN response only keeps 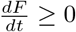 while virus and DVGs rise. We first analyse this model, in which IFN follows viral induction (*sI*) and constant extracellular clearance (*αF* ):

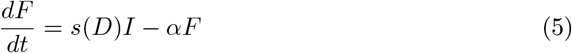

At 24 hours, IFN reaches its peak in the high-DVG group of the *in vivo* data of Penn *et al*. (Fig 3A), the group this argument is about; in the low- and medium-DVG groups IFN peaks at 48 hours instead. At this maximum, production approximately balances clearance:

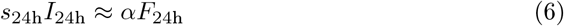

An IFN drop at 48 hours requires 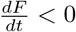. This model keeps 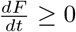:

1. Because *V*_48h_ *> V*_24h_, the infected cell population (*I*_48h_) remains high.
2. The DVG-dependent induction rate is a saturating function: *s*(*D*) = *s*_0_ + *s*_*d*_*D/*(1 + *θD*). This function is strictly increasing in *D*, and *D*_48h_ *> D*_24h_, so *s*_48h_ ≥ *s*_24h_.
3. The total IFN source term at 48 hours is at least as large as it was at 24 hours:

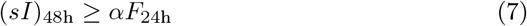
4. Meanwhile, the clearance term at 48 hours is much smaller because the absolute IFN concentration has already fallen (*F*_48h_ ≪ *F*_24h_).

Substituting these facts into the model equation gives:

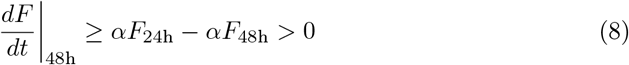

Constant clearance (*α*) keeps 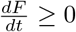 while the viral stimulus stays high.

#### Contradiction: the model with positive IFN response only forces 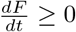 while the data show an IFN drop

This model couples IFN to the viral trajectory. It keeps 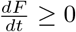 at 48 hours, while the measured IFN in that group drops 8-fold between 24 and 48 hours.

#### Cumulative-IFN negative feedback resolves this contradiction

As IFN accumulates, the factor *g*(cum*F* ) (Eq. 3) lowers the induction source independently of current viral load. Early in infection, cum*F* ≪cum*F*_50_, so *g*(cum*F* ) ≈1 and IFN production proceeds. By 48 hours, the early IFN peak has raised cum*F*, and *g*(cum*F* ) reduces the effective production rate, consistent with negative IFN feedback:

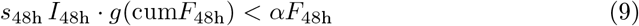

The reduction lets the constant clearance term exceed the induction term, so 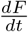 becomes negative at the later time. This reproduces the early IFN decline even while virus and DVGs remain abundant.

### Fitting virus and IFN alone, without the DVG compartment

Most infection experiments measure virus and IFN but not DVGs, even though DVGs arise in almost any influenza infection. We therefore asked what this model recovers from virus and IFN alone. We refit it without the DVG compartment: the *D* state and the parameters *d, h, D*_0_, *s*_*d*_ and *θ* are dropped, the induction term *s*(*D*) *I* reduces to *s*_0_*I*, the virus production term loses its (1 −*d*) factor, and the DVG observations leave the likelihood. Everything else, including the priors, the group sharing and the sampler settings, matches the with-DVG fit. All chains converged, with no divergent transitions.

The cumulative-IFN feedback comes back in the same place and at a similar strength. Its threshold in the high-DVG group is cum*F*_50_ = 30.2 pg day mL^*−*1^ (90% credible interval 9.2–82.7), against 20.5 (5.2–57.8) in the with-DVG fit, and the cumulative exposure reaches 3.1 times that threshold by day 4.5, against 4.3 times with DVGs. In both fits the feedback stays open in the low- and medium-DVG groups, where their thresholds carry a floor (S2 Table). So the inoculum that gives the sharp IFN decline is the one that engages the feedback, whether or not the DVG measurements enter the likelihood.

Both structures reproduce the measured virus and IFN time courses. The DVG compartment earns its place by predicting the DVG measurements themselves and by carrying the mechanism to other DVG doses, rather than by fitting virus and IFN better. Posteriors and the fit figure for the no-DVG model are deposited on GitHub.

### Applications: classic pony and human-titre datasets revisited (no cumulative-IFN feedback)

Both classical datasets analysed here — the equine pony data of Pawelek *et al*. [34] and the human titre data of Baccam *et al*. [40] — predate the cumulative-IFN-exposure formulation introduced in this paper. We apply the

DVG-extended model in its no-cumulative-IFN-feedback variant (*g*(cum*F* ) = 1). For the pony data [34], IFN measurements are present and the IFN equation is retained as *dF/dt* = *s*(*D*) *I* −*αF* . For the human-titre data [40], IFN measurements are absent and *F* is omitted entirely. This isolates virus–DIP competition as a candidate mechanism in the absence of an explicit IFN-shutoff term. The double peak in the human titre data would be worth exploring in its own right. Our fits here reproduce the shape but do not try to separate the mechanisms that could produce it.

#### Pony model (S3 Fig)

The pony fits use the DVG-extended model with a DIP-dependent cell-entry term and no cumulative-IFN feedback. States are *T, I, V, D, F* :

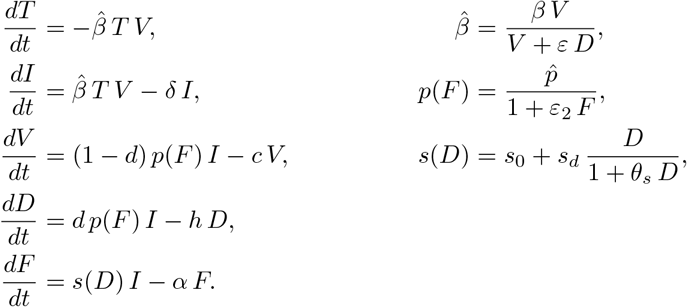

Here *ε* is the virus–DIP competition strength at cell entry and *ε*_2_ is IFN suppression of viral output. Because this model carries no genome-to-particle conversion, *ε* also absorbs the fraction of defective genomes that are not packaged into particles. Both symbols are local to this equine model: elsewhere in the paper *ε* denotes IFN suppression of output and no *ε*_2_ appears.

#### Human-titre model (S4 Fig)

The DIP variants of the Baccam fits use a two-stage eclipse (*I*_1_, *I*_2_) with a DIP-dependent cell-entry term and no IFN state (DIP-only). The DIP-only system is

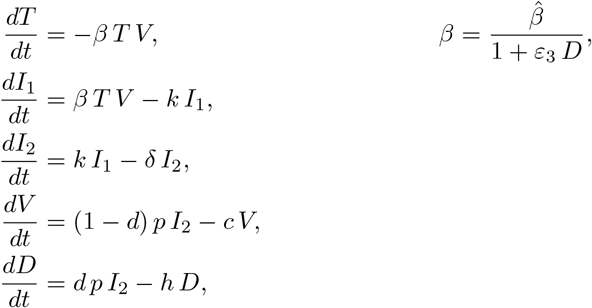

with *k* the eclipse-to-productive transition rate and *ε*_3_ the DIP-load entry-shield strength; initial conditions *I*_1_(0) = *I*_2_(0) = 0, *T* (0) = *T*_0_, *V* (0) = *V*_0_, *D*(0) = *D*_0_. The DIP+IFN variant adds the Baccam IFN term.

#### Earlier or later IFN shutoff changes the host trade-off; no single cum*F*_50_ minimises all four costs at once

After fitting the cumulative-IFN half-saturation threshold (cum*F*_50_), we asked how host outcome would change if this threshold were higher or lower. For each DVG inoculum group, we held all other parameters at their fitted posterior medians and varied only cum*F*_50_: earlier shutoff (cum*F*_50_*/*2), delayed shutoff (10 cum*F*_50_), and feedback removed (*g*(cum*F* ) ≡1). The released case is the dashed curve of Fig 3C. Earlier shutoff lowered the IFN peak and the area under the IFN curve but raised the viral peak and prolonged the high-virus period; delayed shutoff did the reverse, and removing the feedback entirely sat at the far end of this trade-off with the highest IFN and the lowest virus.

The same DVG inoculum group occupied different feedback states depending on cum*F*_50_. No single cum*F*_50_ minimised all four costs (IFN exposure, virus, infected cells, target cells) simultaneously. The preferred shutoff time depends on which cost matters most.

**Fig S1.**
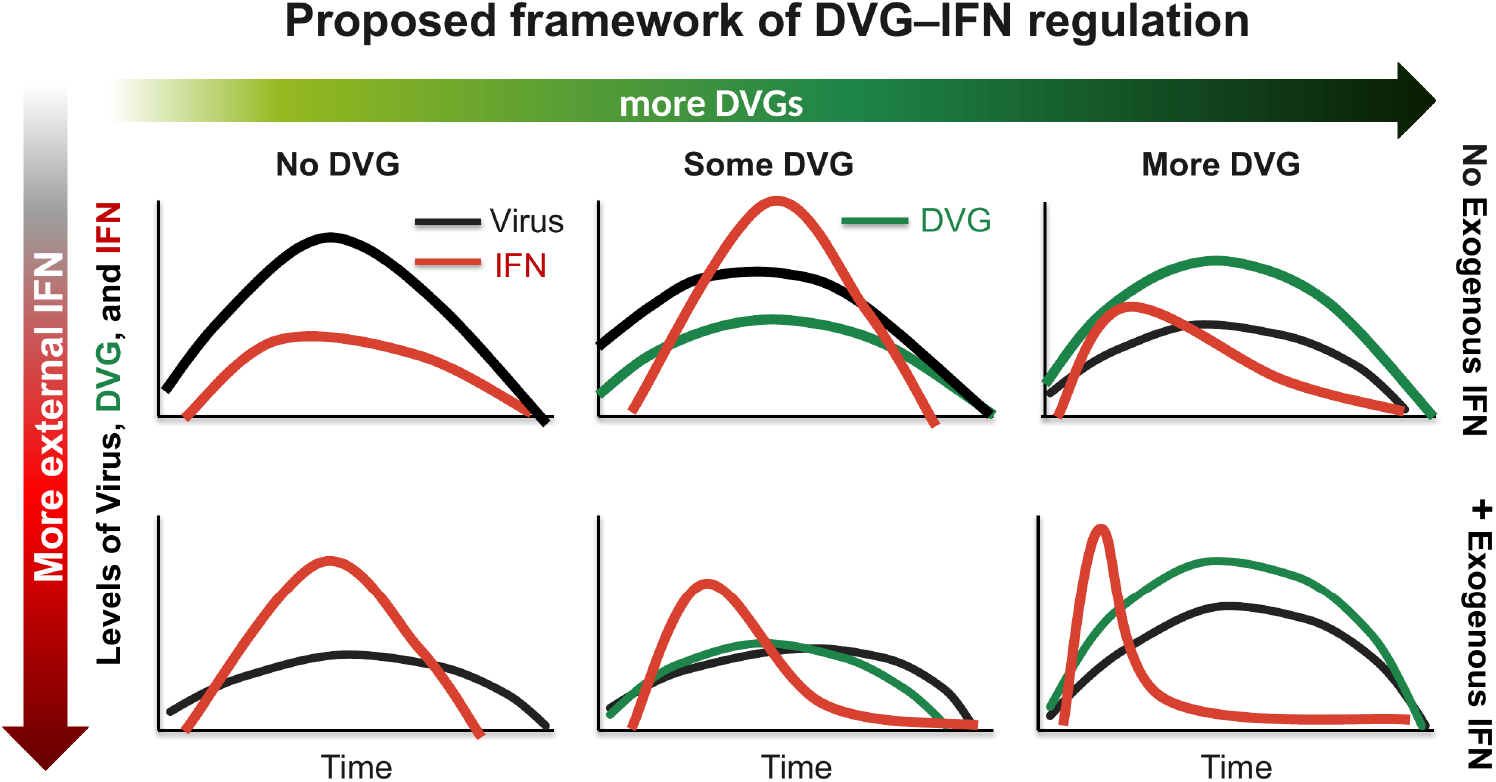
Schematic of virus particle, DVG, and IFN dynamics as a function of DVG and exogenous IFN levels. Columns: no DVG, some DVG, and high DVG levels. Rows: absence (top) and presence (bottom) of exogenous IFN.

**Fig S2.**
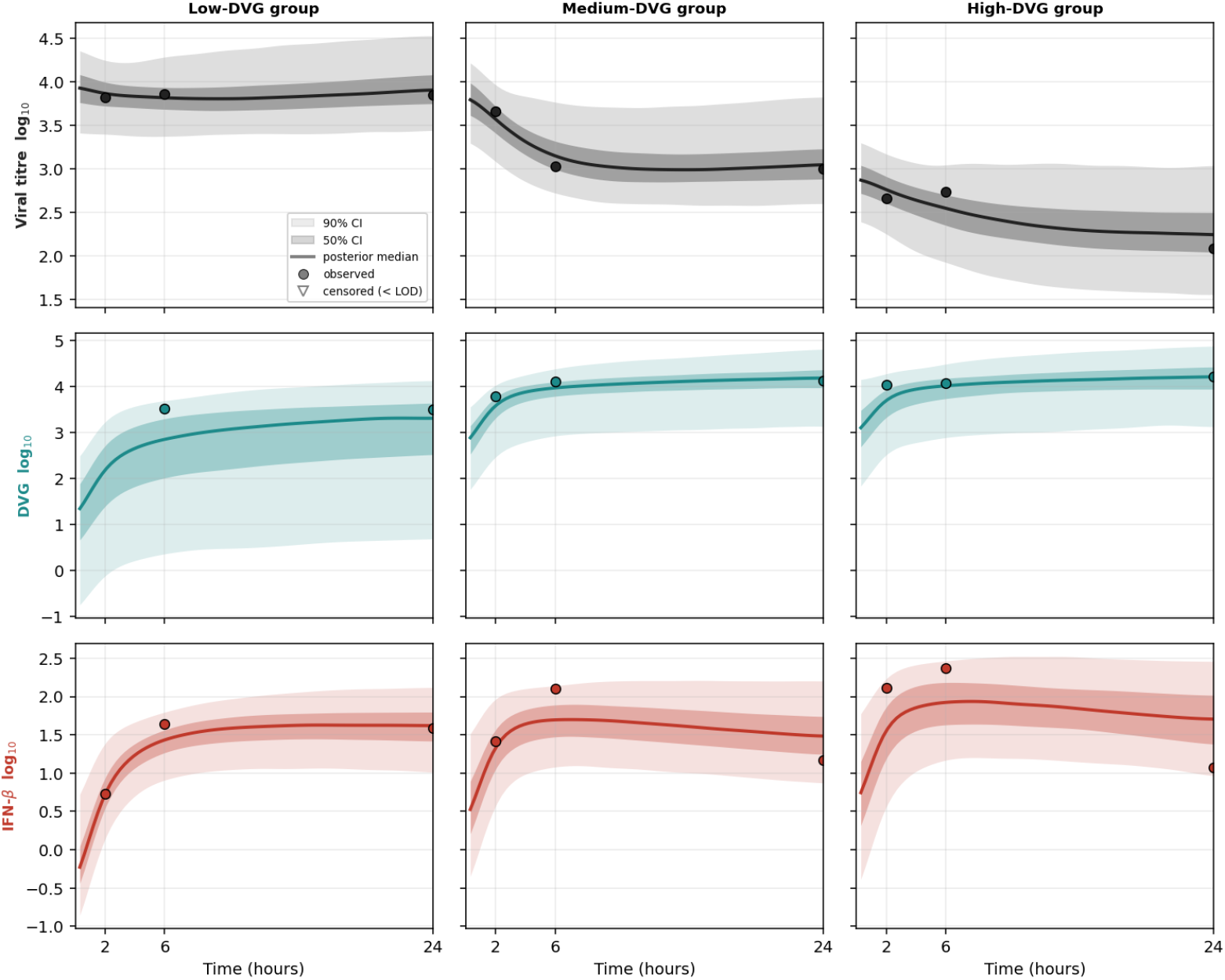
*In vitro* DVG-extended model fits to virus, DVG, and IFN-*β* mRNA time courses. Same data and DVG groups as S5 Fig. The DVG-extended model jointly fits virus (PB1 full genome), DVG (PB1 DVGs), and IFN-*β* mRNA. Rows show virus, DVG, and IFN-*β* mRNA; columns show the low-, medium-, and high-DVG groups, with one *y* axis per row.

**Fig S3.**
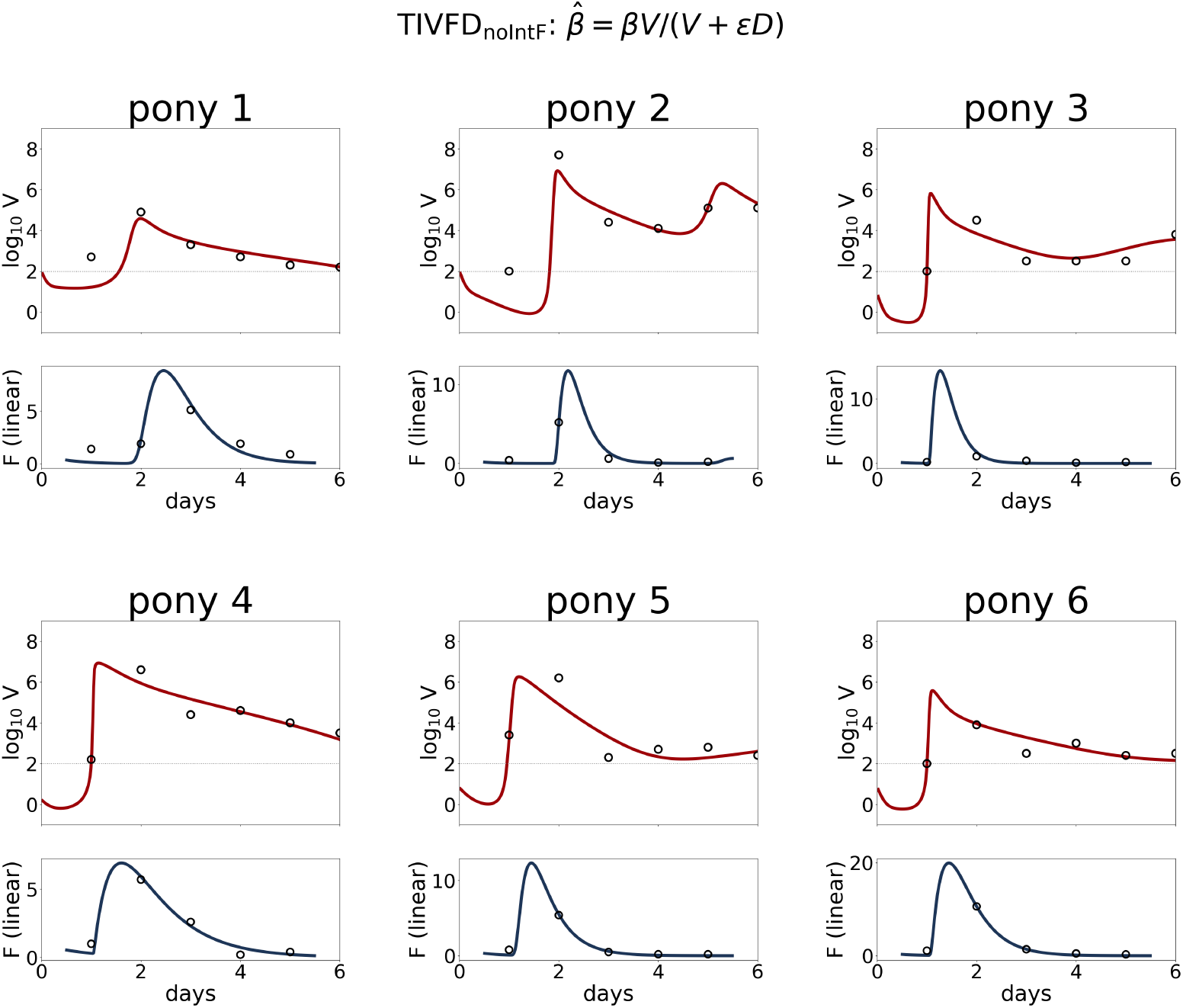
DVG-included fits to the equine influenza pony data. The model was fit to the first 6 days of viral-load and IFN data from the six ponies analysed by Pawelek *et al*. [34]. Each column shows one pony, with viral load on the top row and IFN on the bottom row. Model: the within-host DVG model (Eq 1) with *β* replaced by a DIP-dependent cell-entry term 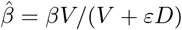 and no cumulative-IFN feedback (*dF/dt* = *s*(*D*) *I − αF* ). Per-pony maximum a posteriori fits; key parameters span *β ≈* 0.7–3 *×* 10^*−*4^, the competition strength *ε ≈* 0.2–4, and the DVG production fraction *d ≈* 0.2–0.5.

**Fig S4.**
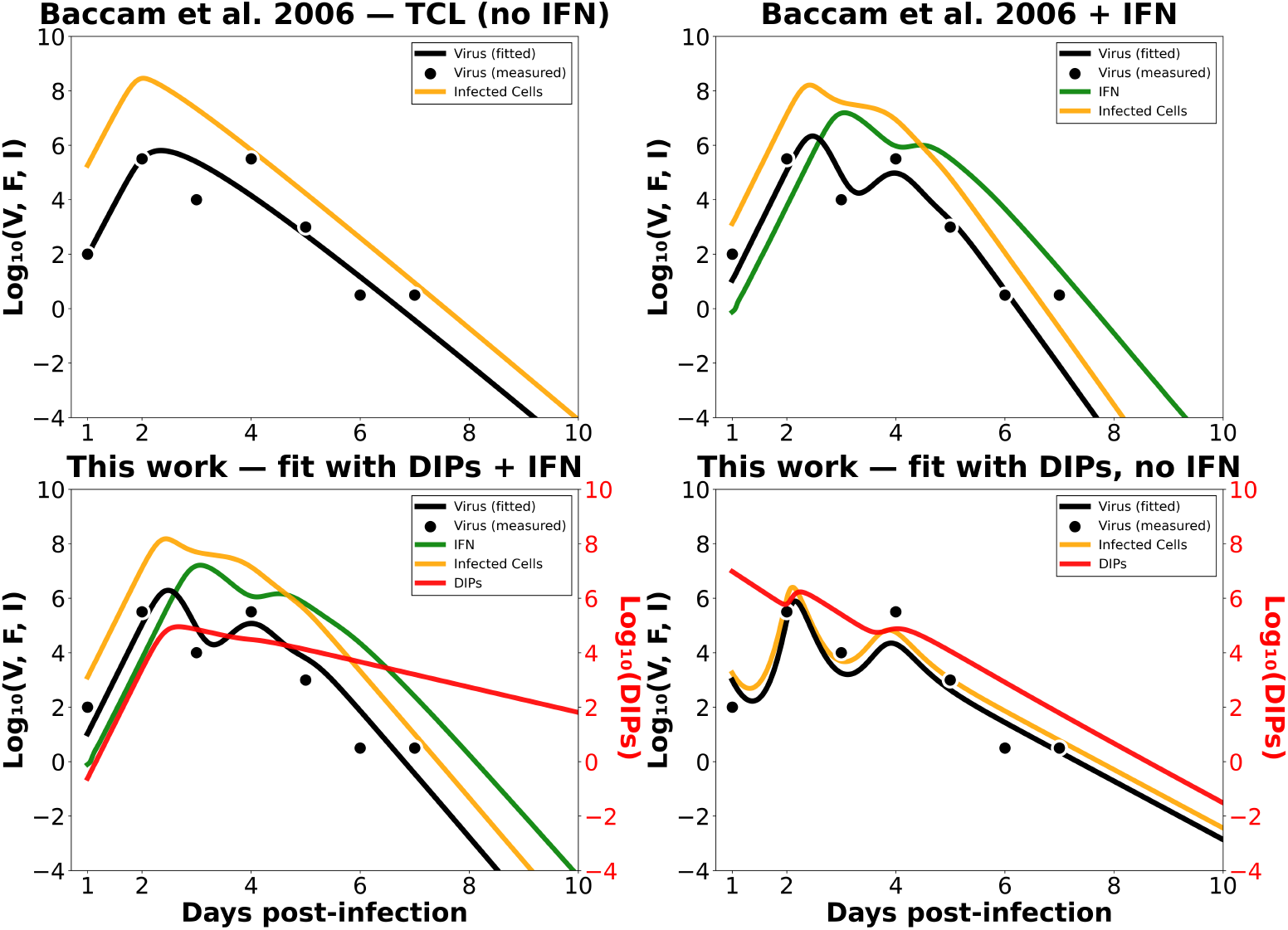
Revisiting classic human influenza viral kinetics with target-cell, IFN, and DIP-based models. The four panels compare model interpretations of the human influenza data analysed by Baccam *et al*. [40]. The target-cell-limited model explains viral decline through target-cell limitation. The Baccam IFN model adds host antiviral control to reproduce non-monotone viral kinetics. The DIP+IFN model includes both host IFN effects and virus–DIP competition. The DIP-only model shows that virus–DIP competition can reproduce the main viral peak and post-peak decline without an explicit IFN variable. The DIP variants use a two-stage eclipse (*I*_1_, *I*_2_) with a DIP-dependent cell-entry term 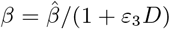 and output partition (1 *− d*); the DIP-only variant omits IFN. DIP-only patient-1 fit: 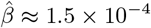, *ε*_3_ *≈* 9.5 *×* 10^*−*6^, *d ≈* 0.30, *p ≈* 20, *c ≈* 30, *δ ≈* 20 (day^*−*1^), *D*_0_ *≈* 292.

**Fig S5.**
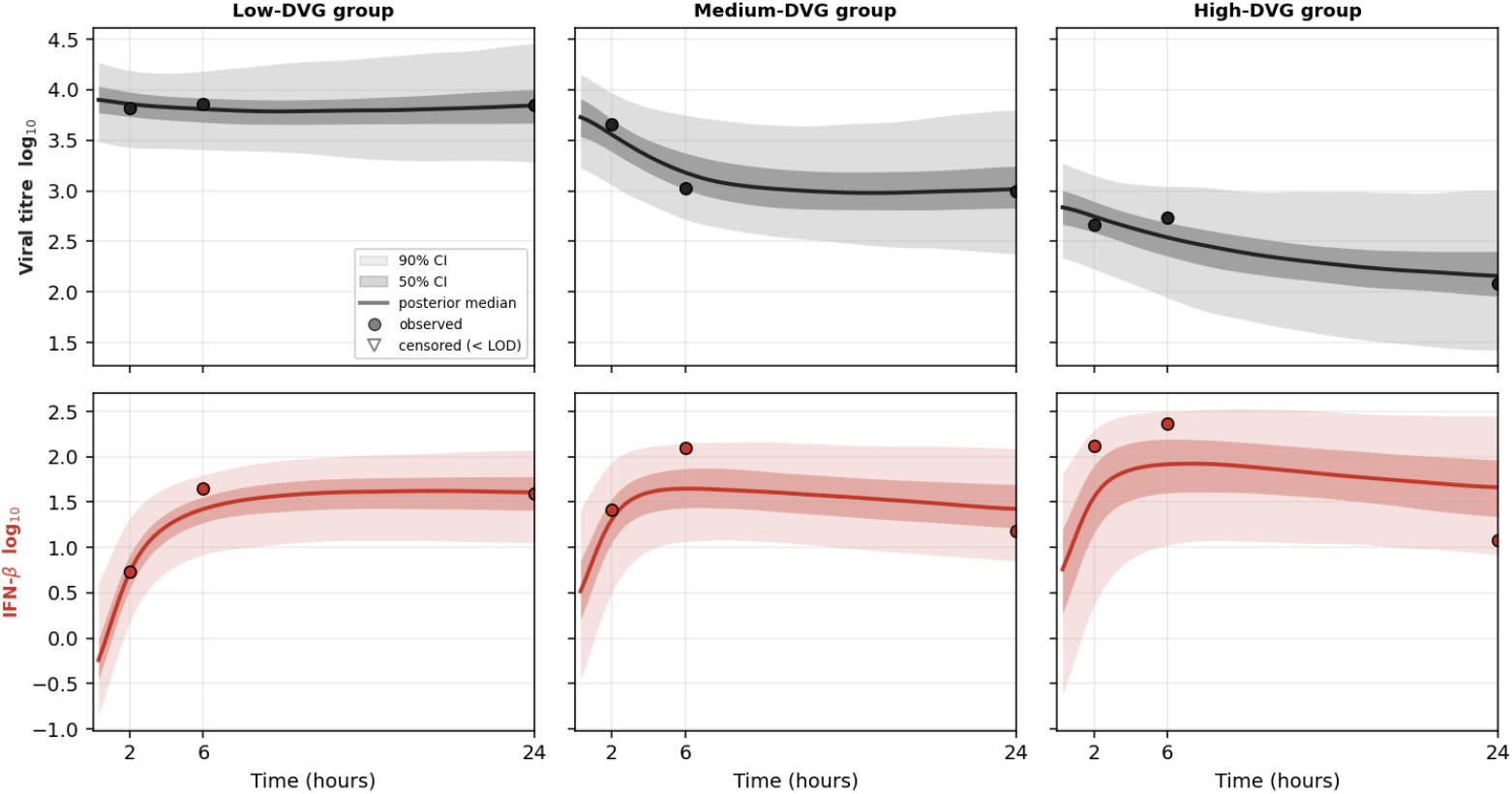
*In vitro* no-DVG model fits to virus and IFN-*β* mRNA time courses. Penn *et al*. [20] PB1 full-genome and IFN-*β* mRNA data, grouped by DVG group (low, med, high). Lines: posterior median. Shaded bands: pointwise posterior quantile intervals (50%: 25%–75%; 90%: 5%–95%). Points: observed data. The no-DVG model fits *V* and *F* only; it does not track DVGs. Rows show virus and IFN-*β* mRNA; columns show the low-, medium-, and high-DVG groups, with one *y* axis per row.

**Fig S6.**
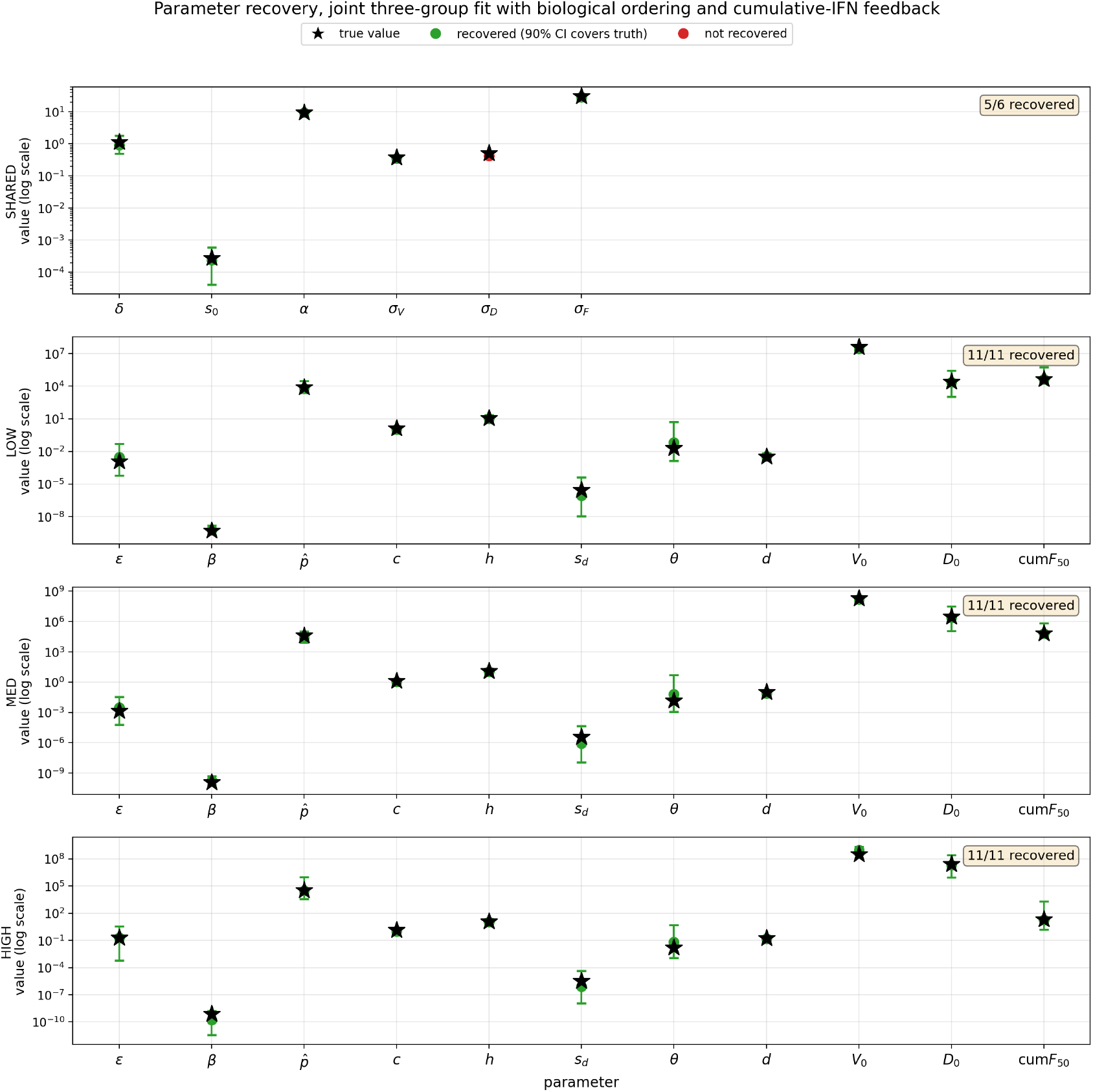
Simulation-based parameter recovery for the with-DVG, cumulative-IFN-feedback model. For each inoculum group (LOW, MED, HIGH), the posterior median of every fitted parameter (S1 Table) was taken as ground truth (black stars). The six-state ODE (Eq 1) was forward-simulated at these values at the Penn *et al*. [20] observation times, and Stan-likelihood-matched noise was added: *V* and *D* Normal on the log_10_ scale, *F* Normal on the linear pg mL^*−*1^ scale it is fitted on. The same Stan model was then refit to the synthetic data under the same priors, bounds, and NUTS settings. Green dots with bars give the posterior median and 90% credible interval of each refit parameter; a parameter counts as recovered when its 90% interval covers the truth. The fit has 39 estimated quantities: six held to one value across the groups (*δ, s*_0_, *α, σ*_*V*_, *σ*_*D*_, *σ*_*F*_ ) and eleven free or partially pooled in each of the three. Thirty-eight of the 39 are recovered, including every rate, the defective-genome fraction *d* and the threshold cum*F*_50_ in all three groups. The remaining one is the DVG observation noise *σ*_*D*_ (red dot), whose truth of 0.514 lies 2% above the upper end of its interval, 0.503.

**Fig S7.**
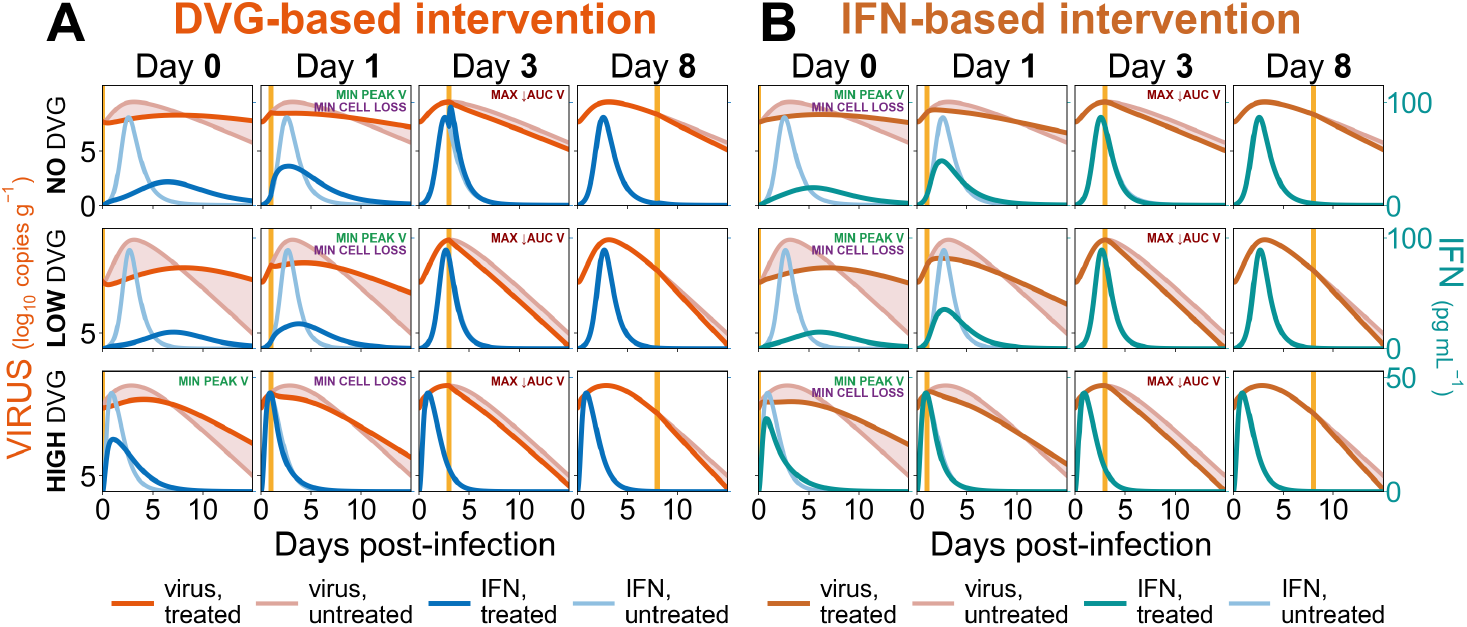
Simulated DVG- and IFN-based interventions across initial DVG levels and treatment times. **(A)** DVG-based intervention and **(B)** IFN-based intervention initiated on days 0, 1, 3, or 8 (yellow lines). Rows show the no-, low-, and high-DVG conditions. Virus and IFN trajectories are shown for treated and untreated conditions; shading indicates the difference in viral trajectories. Curves show the rank-trimmed mean across 200 posterior draws. MIN V indicates the treatment time producing the largest reduction in viral burden within each DVG condition.

#### Footnotes

1 Tentative identification; subject to confirmation with the experimental collaborators.

